# Lipid-Gated Vesicular Trafficking Directs HSPA1A to the Plasma Membrane Through the Endo-Lysosomal Network

**DOI:** 10.64898/2026.06.11.731635

**Authors:** Jensen Low, Azalea Blythe Cuaresma, Clarisse K. Martin, Allen Badolian, Maha AlSebaye, Robert V. Stahelin, Nikolas Nikolaidis

**Affiliations:** Department of Biological Science, Center for Applied Biotechnology Studies, and Titan Supercomputing Center, California State University Fullerton, Fullerton, CA, USA; Borch Department of Medicinal Chemistry and Molecular Pharmacology and The Purdue Institute of Inflammation, Immunology and Infectious Disease, Purdue University, 47907, West Lafayette, IN, USA

**Keywords:** HSPA1A trafficking, Endo-lysosomal pathway, Vesicular transport, Phosphatidylinositol 4-phosphate, Bis(monoacylglycerol)phosphate, Plasma membrane localization, Heat shock response

## Abstract

HSPA1A is a stress-inducible molecular chaperone that localizes to the plasma membrane (PM) of heat-shocked and cancer cells, where its membrane-associated form contributes to therapeutic resistance, membrane stabilization, and immune modulation. Because HSPA1A lacks a signal peptide, it does not follow the classical secretory pathway; instead, it reaches the PM through unconventional routes whose vesicular intermediates and intracellular lipid requirements remain largely undefined. Here, we show that following heat shock, HSPA1A undergoes coordinated redistribution across the endo-lysosomal network. It transiently associates with PI(3)P-enriched early endosomes, progresses through Rab4A- and Rab4B-positive recycling endosomes, and accumulates in LAMP1-positive lysosomes, while avoiding degradative and slow-recycling routes. Pharmacological inhibition of the ER-Golgi pathway did not affect HSPA1A’s PM localization, while disruption of endosomal maturation and lysosomal function resulted in significant reductions. Heat shock drives a progressive increase in lysosomal BMP immunoreactivity, and pharmacological BMP accumulation increased PM-HSPA1A, whereas intracellular antibody-mediated BMP blockade reduced it, identifying BMP-enriched lysosomes as regulatory hubs that govern HSPA1A PM competence. Using a rapamycin-inducible compartment-specific phosphatase system, we further demonstrate that PI(4)P is required not only at the PM for final docking but within early endosomes, late endosomes, and lysosomes, establishing a distributed PI(4)P requirement across the endosomal network. Together, these findings define a lipid-gated vesicular trafficking mechanism for HSPA1A PM localization and identify lysosomal BMP and endosomal PI(4)P as additional regulatory layers relevant to cancer cells in which constitutive lipid remodeling may sustain membrane-associated HSPA1A and its pro-survival functions.

## 1. Introduction

Heat shock protein A1A (HSPA1A) is a 70-kDa stress-inducible molecular chaperone that plays a central role in maintaining protein homeostasis under conditions of cellular stress. Upon induction, HSPA1A stabilizes unfolded proteins, facilitates their refolding, and prevents aggregation, thereby supporting cell survival under adverse conditions ^1,2^. In cancer cells, HSPA1A is frequently overexpressed and contributes to enhanced survival, resistance to apoptosis, and adaptation to the tumor microenvironment ^3,4^. Beyond its well-established cytosolic chaperone functions, HSPA1A also localizes to the plasma membrane (PM) of heat-shocked and tumor cells, where its membrane-associated form (mHSPA1A) is linked to increased therapeutic resistance, membrane stabilization, and immunomodulatory functions, including recognition by natural killer cells and activation of macrophages ^5–8^. Understanding the mechanisms that govern HSPA1A PM translocation is therefore directly relevant to cancer biology and may reveal novel therapeutic vulnerabilities.

Despite its clinical significance, the pathway by which HSPA1A reaches the PM remains poorly understood. HSPA1A lacks a signal peptide, transmembrane domains, and canonical membrane-targeting motifs, and therefore does not follow the classical endoplasmic reticulum (ER)-Golgi secretory route used by most PM-destined proteins ^9,10^. This has led to growing interest in unconventional protein secretion (UPS) pathways as potential mechanisms underlying HSPA1A PM localization. UPS encompasses mechanistically distinct routes by which leaderless cytosolic proteins reach the PM or extracellular space without transiting the ER-Golgi system. Among these, Type II UPS involves vesicle-mediated trafficking through endosomal or lysosomal compartments followed by membrane fusion events, while Type III UPS involves incorporation of cytosolic cargo into late endosomes to form multivesicular bodies (MVBs) that are subsequently trafficked to specific destinations, including the PM ^9,10^. Given HSPA1A’s reported associations with endo-lysosomal compartments and extracellular vesicle pathways, its PM translocation is most consistent with Type II or Type III UPS mechanisms.

Previous studies investigating HSPA1A unconventional secretion reported that extracellular HSPA1A release is largely insensitive to pharmacological disruption of the ER-Golgi pathway, supporting the view that HSPA1A utilizes non-classical trafficking mechanisms ^11–17^. However, these studies primarily measured HSPA1A in the extracellular medium and did not address the route by which HSPA1A becomes associated with the plasma membrane. As a result, the trafficking pathways and molecular determinants governing PM-localized HSPA1A remain unresolved.

A defining feature of HSPA1A membrane association is its dependence on specific lipid interactions. Although HSPA1A lacks canonical lipid-binding domains, multiple studies have demonstrated that it associates selectively with anionic phospholipids, including phosphatidylserine (PS), phosphatidylinositol monophosphates, and bis(monoacylglycerol)phosphate (BMP), a lipid enriched in late endosomal and lysosomal membranes ^16,18–24^. Recent work from our laboratory has established that heat shock triggers a rapid increase in PS levels, which is both necessary and sufficient to drive HSPA1A PM localization, and that PS depletion nearly abolishes heat-induced PM translocation ^25^. In parallel, we demonstrated that phosphatidylinositol 4-phosphate (PI(4)P) accumulates at the PM following heat shock through activation of PI4KIIIα, and that PI(4)P is essential for HSPA1A PM localization, while the closely related PI(4,5)P₂ is not ^26,27^. Together, these findings support a model in which HSPA1A PM translocation is governed by a multi-lipid code in which PS and PI(4)P act as essential docking signals at the cell surface. However, the vesicular route by which HSPA1A reaches the PM and the lipid requirements within endosomal compartments along that route have not been defined.

The endo-lysosomal system ^15–17,19,28,29^ provides a network of interconnected vesicular compartments that regulate cargo sorting, recycling, and degradation, and offers plausible routes for unconventional PM delivery of leaderless proteins. Early endosomes, marked by Rab5 and enriched in PI(3)P, serve as primary sorting stations from which cargo can be directed into fast recycling pathways via Rab4A and Rab4B, slow recycling via Rab11, or degradative pathways via Rab7-positive late endosomes and LAMP1-positive lysosomes ^30–33^. Lysosomes, in addition to their degradative role, can undergo regulated exocytosis and fuse with the PM under stress conditions, making them candidate intermediates for unconventional PM delivery ^34,35^. PI(4)P is known to regulate membrane identity and vesicular trafficking specificity across multiple compartments of this network, and its compartment-specific dynamics are controlled by lipid kinases and phosphatases including PI4KIIIα and Sac1 ^36–42^. Whether PI(4)P within endosomal compartments, rather than solely at the PM, contributes to HSPA1A trafficking has not previously been examined.

Here, we investigate the vesicular route and lipid determinants of HSPA1A trafficking to the PM following heat shock. Using compartment-specific Rab markers, pharmacological perturbation of vesicular trafficking pathways, modulation of lysosomal lipid composition, and a rapamycin-inducible system for compartment-targeted PI(4)P depletion, we tested the hypothesis that HSPA1A is trafficked to the PM through a lipid-dependent vesicular mechanism involving the endo-lysosomal network, in which PI(4)P is required across multiple endosomal compartments, and BMP-enriched lysosomal compartments serve as regulatory hubs for PM localization competence.

## 2. Materials and Methods

### 2.1 Cell Line and Culture

HeLa cells (ATCC® CCL-2™), derived from Henrietta Lacks, were used for all experiments. Cells were maintained in Minimum Essential Medium (MEM) supplemented with 10% fetal bovine serum, 2 mM L-glutamine, 0.1 mM non-essential amino acids, 1 mM sodium pyruvate, and penicillin-streptomycin. Cultures were maintained at 37°C in a humidified atmosphere containing 5% CO₂. Cell viability was assessed after all treatments using a trypan blue exclusion assay and quantified with the Cellometer® Auto X4 Cell Counter (Nexcelom Bioscience, Manchester, UK). No major reduction in viability was observed across conditions.

### 2.2 Plasmids and Constructs

To visualize HSPA1A subcellular localization, fluorescently tagged constructs were used as previously described ^25–27,43^. Briefly, the mouse *hspa1a* cDNA sequence (accession number BC054782; Open Biosystems, GE Dharmacon) was subcloned into pEGFP-C2 or mRFP-C1 vectors to generate N-terminally tagged HSPA1A-GFP and HSPA1A-RFP, respectively.

To assess colocalization with endosomal and lysosomal compartments, the following constructs were used: EGFP-Rab4A (Addgene plasmid #49434; gift from Marci Scidmore) ^44^, EGFP-Rab4B (Addgene plasmid #49468; gift from Marci Scidmore) ^44^, Rab5-mCherry (Addgene plasmid #49201; gift from Gia Voeltz) ^45^, DsRed-Rab7-WT (Addgene plasmid #12661; gift from Richard Pagano) ^46^, Rab11-DsRed (Addgene plasmid #12679; gift from Richard Pagano) ^46^, GFP-EEA1 wild type (Addgene plasmid #42307; gift from Silvia Corvera) ^47^, GFP-hPIKfyve (Addgene plasmid #121148; gift from Geert van den Bogaart) ^48^, pDEST-CMV 3xFLAG-LC3A-GFP (Addgene plasmid #123106; gift from Robin Ketteler) ^49^, and LAMP1-RFP (Addgene plasmid #1817; gift from Walther Mothes) ^50^.

To test whether PI(4)P within specific endosomal compartments is required for HSPA1A PM localization, a rapamycin-inducible dimerization system was used to recruit the Sac1 phosphatase to defined compartments. This system uses FKBP-tagged phosphatase constructs and FRB-tagged compartment anchors, which dimerize upon rapamycin addition, targeting enzymatic activity to a specific membrane. Constructs used were: pJ-Sac1-FKBP-mRFP (Sac1-WT; Addgene plasmid #38000; gift from Robin Irvine) ^51^, pJ-Dead-FKBP-mRFP (catalytically inactive control; Addgene plasmid #38002; gift from Robin Irvine) ^51^, pJ-WT-FKBP-mRFP (Addgene plasmid #37999; gift from Robin Irvine) ^51^. To direct Sac1 recruitment to early and late endosomes, iRFP-FRB-Rab5 (Addgene plasmid #51612; gift from Tamas Balla) ^52^ and iRFP-FRB-Rab7 (Addgene plasmid #51613; gift from Tamas Balla) ^52^ were used as compartment-specific FRB anchors, respectively. To assess PI(4)P requirements in lysosomal compartments specifically, pLJM1-Lyso-FLAG-GFP-Sac1-Cat-WT (Addgene plasmid #134645; gift from Roberto Zoncu) ^53^ was used as the active lysosomal-targeted phosphatase, with pLJM1-Lyso-FLAG-GFP-Sac1-Cat-CS (Addgene plasmid #134653; gift from Roberto Zoncu) ^53^ serving as the catalytically inactive control.

Recombinant plasmids were transformed into DH5α *E. coli* and selected on appropriate antibiotic plates. Plasmids were purified using ZymoPURE™ Miniprep or Midiprep Kits (Zymo Research, Irvine, CA) according to the manufacturer’s instructions.

### 2.3 Cell Transfections

To express fluorescently tagged HSPA1A and subcellular marker constructs, cells were transiently transfected using PolyJet™ In Vitro DNA Transfection Reagent (SignaGen Laboratories, Rockville, MD) according to the manufacturer’s instructions. One day before transfection, HeLa cells were seeded onto poly-D-lysine (PDL)-coated glass coverslips in 24-well plates at 2.0 × 10⁴ cells per well. After 18 hours, cells were transfected with 1 µg total DNA per well. For co-transfections involving two plasmids, 0.5 µg of each was used; for three plasmids, approximately 0.33 µg per plasmid. Transfected cells were incubated for at least 18 hours prior to heat shock or pharmacological treatment.

### 2.4 Heat Shock Protocol

To induce cellular stress and trigger HSPA1A redistribution, cells were subjected to heat shock at 42°C for 1 hour in a humidified CO₂ incubator, followed by recovery at 37°C. Two recovery timepoints were examined: 0 hours (cells fixed immediately after heat shock) and 8 hours (cells returned to 37°C for 8 hours post-stress). Control cells were maintained at 37°C throughout. Heat shock and pharmacological treatment timelines were coordinated so that both concluded simultaneously at the time of fixation.

### 2.5 Pharmacological Treatments

To test whether specific vesicular trafficking pathways or lipid species contribute to HSPA1A PM localization, cells were treated with pharmacological agents targeting distinct cellular processes. Compounds were selected based on established mechanisms of action and grouped accordingly. All treatments were applied during or after transfection, with concentrations and exposure times optimized based on published studies. A summary of all pharmacological treatments is provided in Table 1.

**Table 1.** Pharmacological treatments used to modulate vesicular trafficking and lipid composition.

| Category | Compound | Concentration | Exposure time |
| --- | --- | --- | --- |
| ER-Golgi inhibitors | Brefeldin A | 10 μM | 9 hours |
| | Exo1 | 50 μM | 9 hours |
| | Exo2 | 50 μM | 9 hours |
| | Tunicamycin | 20 μM | 6 hours |
| Endosomal/lysosomal inhibitors | EGA | 20 μM | 9 hours |
|  | Bafilomycin A1 | 100 nM | 6 hours |
| BMP modulators* | Amiodarone | 10 μM | 24 hours |
| | Fosinopril | 10 μM | 24 hours |
| Dimerization inducer | Rapamycin | 1 μM | 7 minutes |

\*For dose-response validation experiments, Amiodarone and Fosinopril were applied at 0.1, 0.5, 1, 10, and 20 μM for 24h at 37°C. BMP immunoreactivity was assessed by ICC as described in section 2.9.

DMSO was used as a vehicle control for all compounds dissolved in DMSO; methanol was used as a control for Amiodarone and Fosinopril; BioPORTER delivering a mouse IgG isotype control antibody (Invitrogen, catalog #10400C) served as a control for antibody delivery experiments.

### 2.6 Antibody-Mediated BMP Inhibition

To determine whether BMP is functionally required for HSPA1A PM localization, an anti-BMP antibody (clone 6C4, IgG1κ; catalog #Z-PLBPA; Echelon Biosciences, Salt Lake City, UT) was delivered intracellularly using BioPORTER protein delivery reagent (QuikEase Protein Delivery Kit; Thermo Fisher Scientific, Waltham, MA) according to the manufacturer’s instructions. As a control, a mouse IgG isotype control antibody (mouse IgG; catalog #10400C; Invitrogen, Carlsbad, CA) was delivered using BioPORTER under identical conditions. To further validate the intracellular antibody delivery approach, a phosphatidylserine (PS)-specific monoclonal antibody (anti-phosphatidylserine, clone 1H6; catalog #05-719; Sigma-Aldrich, St. Louis, MO) was delivered intracellularly using BioPORTER protein delivery reagent under identical conditions. This treatment served as a positive control based on the established requirement of PS for HSPA1A plasma membrane localization ^25^.

### 2.7 Compartment-Specific Phosphatase Recruitment

To test whether PI(4)P within early endosomes, late endosomes, or lysosomes is required for HSPA1A PM localization, Sac1 phosphatase was recruited to specific compartments using a rapamycin-inducible FKBP-FRB dimerization system. HeLa cells were co-transfected with HSPA1A-GFP, a compartment-specific FRB anchor (iRFP-FRB-Rab5 for early endosomes, iRFP-FRB-Rab7 for late endosomes, or a lysosomal-targeted FRB construct), and either Sac1-WT-FKBP-mRFP or the catalytically inactive Sac1-Dead-FKBP-mRFP. Following heat shock and the indicated recovery period, compartment-specific Sac1 recruitment was induced by treating cells with 1 µM rapamycin for 7 minutes immediately before fixation. Control cells were maintained without rapamycin. The Sac1-Dead construct served as a critical specificity control, confirming that any observed reduction in HSPA1A PM localization reflects phosphatase activity rather than overexpression or targeting artifacts. Following treatment, cells were washed three times with 1× PBS before fixation.

### 2.8 Immunofluorescence and Confocal Microscopy

To visualize HSPA1A subcellular distribution and its colocalization with endosomal and lysosomal markers, cells were fixed and imaged by confocal microscopy following all treatments. After transfection and treatment, cells were fixed at room temperature with 4% paraformaldehyde (PFA) in complete growth medium for 12 minutes, followed by three washes with 1× PBS. To label the plasma membrane, cells were incubated with 1 µg/mL wheat germ agglutinin (WGA) Alexa Fluor® 555 conjugate (Thermo Fisher Scientific, Waltham, MA) for 10 minutes at room temperature, then washed three times with 1× PBS. Coverslips were mounted onto glass slides using DAPI Fluoromount-G® (SouthernBiotech, Birmingham, AL), dried for 24 hours at room temperature in the dark, and stored at 4°C until imaging. In experiments using WGA-350 (the number refers to the fluorophore conjugate) for PM labeling, Fluoromount-G without DAPI was used to avoid spectral overlap.

Cells were imaged using an Olympus FLUOVIEW FV3000 confocal microscope equipped with a 60×/1.42 NA oil immersion objective. For multichannel acquisition, images were collected sequentially using the following settings: green channel (excitation 488 nm, emission 510 nm), blue channel (excitation 405 nm, emission 461 nm), and red channel (excitation 561 nm, emission 583 nm). For colocalization analysis, Z-stack images were acquired with 10 optical sections per cell, distributed at equal intervals between the top and bottom of the cell.

### 2.9 Image Analysis

To quantify HSPA1A PM localization and colocalization with vesicular markers, fluorescence intensity was analyzed using ImageJ ^54^. For Z-stack acquisitions, 10 optical sections were collected per cell, with the top and bottom defined as the first and last sections, respectively, and the intermediate sections distributed at equal intervals by the microscope system.

To quantify colocalization between HSPA1A and compartment-specific markers, three channels were acquired per cell: DAPI, HSPA1A, and the respective marker construct. Analysis was performed on a per-cell basis. Total HSPA1A fluorescence within the cytosol, excluding the nucleus, was measured as the denominator. To determine the fraction of HSPA1A associated with marker-positive regions, an ROI corresponding to the marker signal was defined within the same cell and overlaid onto the HSPA1A channel at the identical spatial position. The HSPA1A fluorescence within this ROI served as the numerator. Background correction was applied to both measurements using the corrected total cell fluorescence (CTCF) formula: CTCF = Integrated Density − (Area of selected region × Mean background fluorescence), following established protocols ^25–27,43^. Colocalization was expressed as the CTCF-corrected HSPA1A signal within marker-defined regions relative to total cellular HSPA1A fluorescence.

For PM localization experiments, the PM was identified using the WGA-FA555 channel as a reference, and an ROI was manually traced along the inner boundary of the PM on the HSPA1A channel, isolating signal confined between the outer and inner membrane boundaries, as previously described ^25–27,43^. Total HSPA1A fluorescence within the cytosol, excluding the nucleus, served as the denominator. Background correction was applied using the CTCF formula as described above. PM localization was expressed as the CTCF-corrected HSPA1A signal at the PM relative to total cellular HSPA1A fluorescence. All data represent three independent experiments with n > 30 cells per condition, unless otherwise stated.

To assess intracellular BMP levels, cells were processed for immunocytochemistry (ICC) using the anti-BMP antibody (clone 6C4; catalog #Z-PLBPA; Echelon Biosciences, Salt Lake City, UT) ^16,55^ at 1:400 dilution, followed by Alexa Fluor 555-conjugated secondary antibody at 1:400 dilution (Goat anti-Mouse IgG (H+L) Cross-Adsorbed Secondary Antibody, Alexa Fluor™ 555 - A-21422; Invitrogen, Carlsbad, CA). Briefly, cells were fixed in 4% paraformaldehyde for 15 min at room temperature, washed five times with PBS, and permeabilized with 0.1% NP-40 for 30 min at room temperature. Cells were blocked for 30 min in a blocking solution containing 1× PBS, 10% NGS, and 0.01% Triton X-100. Primary antibody was applied overnight at 4°C, and secondary antibody was applied for 1h at room temperature in the dark, both in blocking solution, with five PBS washes between steps. To quantify intracellular BMP immunoreactivity following heat shock, fluorescence intensity was measured in ImageJ per cell. Two channels were acquired per cell: DAPI and anti-BMP. An ROI corresponding to the cell body was defined by tracing the cell boundary based on the anti-BMP signal, with the nucleus excluded using the DAPI channel. Total anti-BMP fluorescence within this ROI was background-corrected using the CTCF formula as described above and normalized to cell area (μm²). Data are expressed as CTCF-corrected anti-BMP signal per unit cell area [CTCF (anti-BMP/cell area)].

For drug validation experiments, Amiodarone and Fosinopril were applied at concentrations of 0.1, 0.5, 1, 10, and 20 μM for 24h at 37°C before fixation and ICC processing as described above.

### 2.10 Subcellular Fractionation and Western Blotting

To assess the redistribution of HSPA1A across organelle compartments following heat shock, subcellular fractionation was performed at 37°C, 0h, 8h, and 24h recovery timepoints. PM-enriched fractions were isolated using the Minute™ Plasma Membrane Protein Isolation and Cell Fractionation Kit (catalog #SM-005; Invent Biotechnologies, Plymouth, MN) as previously described (Low et al., 2025). Early endosomal enriched fractions were isolated using the Minute™ Endosome Isolation and Cell Fractionation Kit (catalog #ED-028; Invent Biotechnologies, Plymouth, MN). Lysosomal-enriched fractions were isolated using the Minute™ Lysosome Isolation Kit (catalog #LY-034; Invent Biotechnologies, Plymouth, MN). All kits were used according to the manufacturer’s instructions.

Fractions were analyzed by SDS-PAGE and western blotting using the following primary antibodies: anti-HSPA1A (clone C92F3A-5, mouse IgG, 1:1000; Enzo Life Sciences, Farmingdale, NY), anti-Na⁺/K⁺-ATPase α1 (ATP1A1; clone EP1845Y, catalog #2047-1, rabbit RabMAb®, 1:1000; Thermo Fisher Scientific, St. Louis, MO), anti-EEA1 (clone C45B10, rabbit monoclonal, catalog #3288, 1:1000; Cell Signaling Technology, Danvers, MA), anti-LAMP1 (clone D2D11, rabbit monoclonal XP®, catalog #9091, 1:1000; Cell Signaling Technology, Danvers, MA), and anti-GAPDH (clone 1E6D9, catalog #60004-1-Ig, mouse IgG, 1:1000; Proteintech Group, Rosemont, IL). Proteins were transferred to nitrocellulose membranes, blocked in 5% nonfat dry milk in TBST for 1h at room temperature, and incubated with primary antibodies overnight at 4°C. Membranes were then washed in TBST and incubated with horseradish peroxidase-conjugated secondary antibodies for 1h at room temperature. Bound antibodies were visualized using an enhanced chemiluminescence substrate. Signals were detected using the Omega Lum™ C Imaging System (Gel Company, Aplegen, San Francisco, CA) and quantified by densitometry using Image Studio Lite (LI-COR Biosciences, Lincoln, NE).

### 2.11 Statistical Analysis

To determine whether observed differences in HSPA1A PM localization or vesicular Colocalization were statistically significant, one-way analysis of variance (ANOVA) followed by Tukey’s honestly significant difference (HSD) post-hoc test was applied to all datasets. A p-value < 0.05 was considered statistically significant. Adjusted p-values are reported in all figures (* p<0.05, ** p<0.01, *** p<0.001, **** p<0.0001; comparisons not shown were not significant unless otherwise indicated in the text). Box plots were generated using BoxPlotR ^56^, and show the median (center line), interquartile range (box), and whiskers extending 1.5× the interquartile range; crosses indicate sample means.

## 3. Results

### 3.1 Heat Shock Induces Coordinated Redistribution of HSPA1A Across Endosomal Compartments

To determine how HSPA1A redistributes among intracellular vesicular compartments following heat shock, HeLa cells co-expressing HSPA1A-GFP or HSPA1A-RFP with fluorescently tagged compartment-specific markers were subjected to heat shock and analyzed by confocal microscopy at 0h and 8h recovery (Fig. 1 and Supplemental Fig. S1). Colocalization was quantified as the fraction of HSPA1A signal within marker-positive puncta relative to total cellular HSPA1A.

**Fig. 1.**
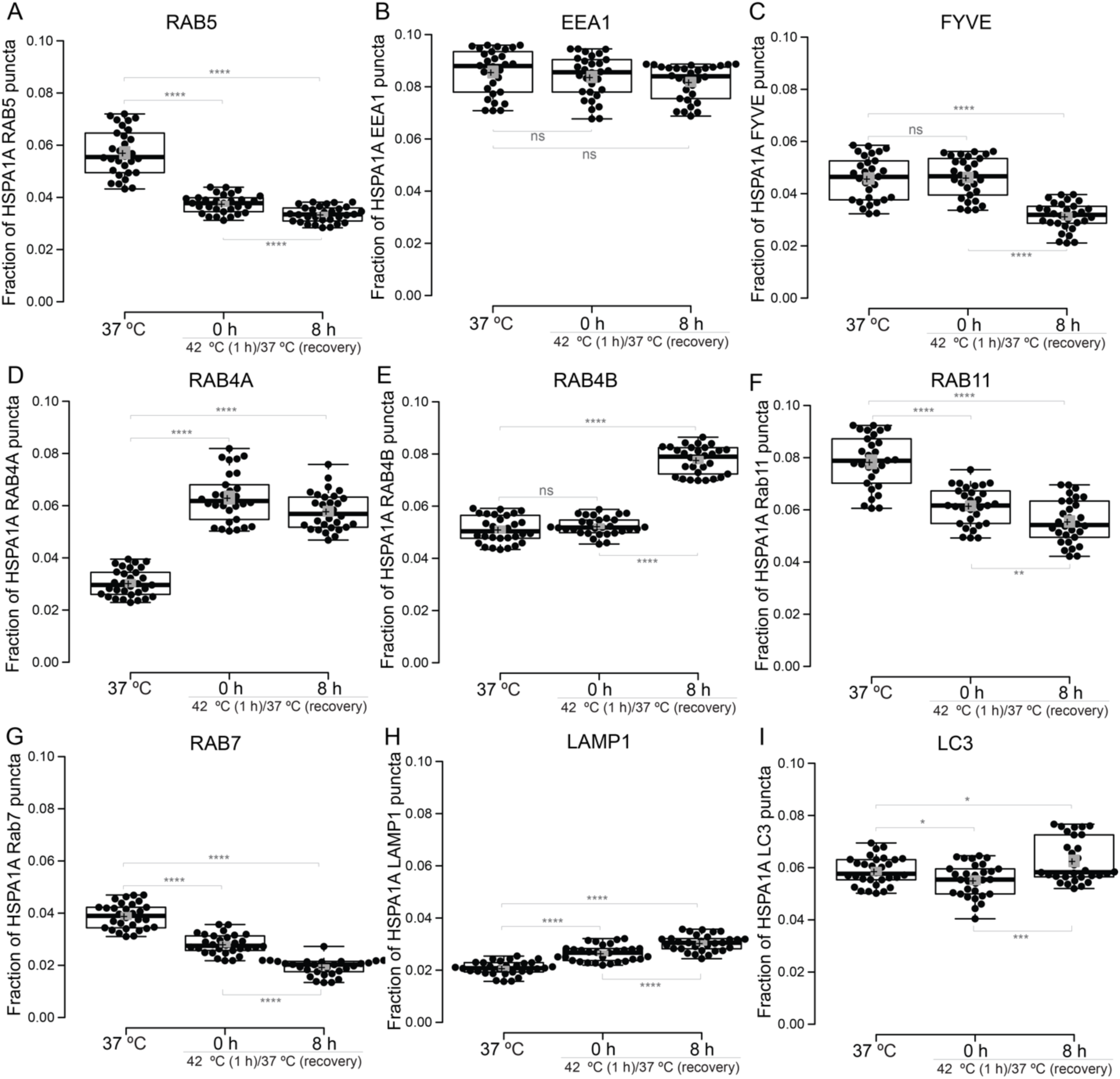
Heat shock induces coordinated redistribution of HSPA1A across endosomal and lysosomal compartments. HeLa cells co-expressing HSPA1A-GFP or HSPA1A-RFP with fluorescently tagged compartment markers were maintained at 37°C or subjected to heat shock (42°C, 1h) followed by recovery at 37°C for 0h or 8h. Colocalization was quantified as the fraction of HSPA1A signal within marker-positive puncta relative to total cellular HSPA1A fluorescence, as described in Materials and Methods. Each data point represents one cell. Boxes show interquartile range with median; whiskers extend to 1.5× IQR. **(A)** Rab5 (early endosome GTPase). **(B)** EEA1 (early endosome tethering factor). **(C)** FYVE (PI(3)P sensor). **(D)** Rab4A (fast recycling endosomes). **(E)** Rab4B (fast recycling endosomes, late phase). **(F)** RAB11 (slow recycling endosomes). **(G)** RAB7 (late endosomes). **(H)** LAMP1 (lysosomes). **(I)** LC3 (autophagosomal membranes). Statistical comparisons by one-way ANOVA with Tukey’s post-hoc test. * p<0.05, ** p<0.01, *** p<0.001, **** p<0.0001. Comparisons not indicated were not statistically significant. N>30 cells per condition from three independent experiments. Representative confocal images for all markers are shown in Supplemental Fig. S1.

To assess whether HSPA1A associates with classical early endosomal compartments following heat shock, Colocalization with Rab5 (early endosome GTPase), EEA1 (early endosome tethering factor), and FYVE [PI(3)P sensor marking early endosomal membranes] was examined. HSPA1A colocalization with Rab5 decreased significantly at both 0h (34.2% reduction vs. 37°C, p<0.0001) and 8h recovery (41.4% reduction, p<0.0001), with a further significant decline between timepoints (10.9%, p<0.0001), indicating progressive departure from classical early endosomes following stress (Fig.1). EEA1 colocalization remained unchanged across all conditions (ANOVA p=0.17), confirming that the canonical early endosome tethering compartment is not a site of HSPA1A retention. FYVE colocalization with HSPA1A was unchanged at 0h compared to baseline (p=0.86) and decreased significantly by 8h recovery (31.7% reduction vs. 37°C, p<0.0001; 32.2% reduction vs. 0h, p<0.0001) (Fig.1). The absence of increased FYVE-HSPA1A colocalization, despite the established functional importance of PI(3)P for HSPA1A PM localization ^27^, is consistent with a competition model in which the FYVE domain and HSPA1A bind to the same PI(3)P-enriched early endosomal membranes. Under conditions where FYVE occupies PI(3)P sites, HSPA1A is displaced, as demonstrated by the approximately 50% reduction in PM localization observed when PI(3)P availability is reduced by wortmannin treatment or masked by EEA1 overexpression ^27^. Together, these data suggest that HSPA1A transiently associates with PI(3)P-enriched early endosomal compartments during or immediately after heat shock, before progressing to downstream recycling and lysosomal routes (Fig. 1 and Supplemental Fig. S1).

To determine whether HSPA1A enters recycling pathways following heat shock, colocalization with Rab4A and Rab4B (fast recycling endosomes) and Rab11 (slow recycling endosome compartment) was examined ^57^. HSPA1A colocalization with Rab4A increased dramatically at 0h recovery (108.0% increase vs. 37°C, p<0.0001) and remained significantly elevated at 8h (90.7% increase vs. 37°C, p<0.0001), with a modest decline between timepoints (8.3%, p=0.018), indicating immediate and sustained engagement of fast recycling endosomes that peaks at the earliest measured timepoint after heat shock (Fig. 1). In striking contrast, Rab4B colocalization showed no significant change at 0h (p=0.26) but increased substantially by 8h recovery (52.2% increase vs. 37°C, p<0.0001; 48.5% increase vs. 0h, p<0.0001), demonstrating a clear temporal shift from Rab4A- to Rab4B-associated compartments during recovery. Rab11 colocalization decreased significantly at both 0h (21.5% reduction, p<0.0001) and 8h (29.2% reduction, p<0.0001), with a further decline between timepoints (9.8%, p=0.004), indicating that slow recycling pathways are actively disengaged following heat shock (Fig. 1).

To assess HSPA1A association with degradative and lysosomal compartments, Colocalization with Rab7 (late endosomes) and LAMP1 (lysosomes) was examined. Rab7 colocalization decreased significantly and progressively, 27.8% at 0h (p<0.0001) and 50.3% at 8h (p<0.0001) compared to 37°C, with a further significant decline between timepoints (31.2%, p<0.0001), indicating that HSPA1A actively avoids the canonical late endosomal degradative route following stress. In contrast, LAMP1 colocalization increased progressively and significantly at both 0h (27.7% increase, p<0.0001) and 8h (46.8% increase, p<0.0001), with continued accumulation between timepoints (15.0%, p<0.0001), indicating sustained association with lysosomal-associated compartments throughout the recovery period (Fig. 1).

To determine whether HSPA1A associates with autophagic vesicles following heat shock, colocalization with LC3, a marker of autophagosomal membranes, was examined. LC3 colocalization showed a small but significant decrease at 0h (-6.0%, p=0.026) followed by a small but significant increase at 8h compared to both baseline (+6.7%, p=0.036) and 0h (+13.5%, p=0.0003), indicating dynamic and temporally regulated engagement of autophagy-linked compartments during the later recovery phase (Fig. 1).

Together, these findings demonstrate that heat shock triggers a coordinated and selective redistribution of HSPA1A across the endo-lysosomal network. The data reveal a clear temporal pattern: immediate and dramatic engagement of Rab4A fast recycling endosomes (>2-fold increase at 0h), a delayed shift to Rab4B compartments (>50% increase at 8h), progressive avoidance of Rab5 classical early endosomes, Rab7 late endosomes, and Rab11 slow recycling routes, transient PI(3)P ^27^ endosomal involvement captured through competition dynamics with the FYVE biosensor, and sustained accumulation in LAMP1-positive lysosomal compartments. This pattern is inconsistent with passive diffusion or nonspecific membrane association and instead reflects active, temporally regulated sorting into specific vesicular pathways consistent with directed unconventional trafficking toward the plasma membrane.

### 3.2 HSPA1A Plasma Membrane Localization Does Not Depend on the ER-Golgi Secretory Pathway

To determine whether HSPA1A reaches the plasma membrane via the classical ER-Golgi secretory pathway, cells were treated with pharmacological inhibitors targeting distinct steps of the conventional protein secretion pathway. Because HSPA1A lacks a signal peptide and canonical membrane-targeting motifs, and based on several reports ^11–17^, we predicted that its PM localization would be insensitive to disruption of this pathway. If correct, this would place HSPA1A within the framework of unconventional protein membrane localization.

The absence of any effect on HSPA1A PM localization across four inhibitors that each block a different step of the classical secretory pathway, BFA ^58^, which collapses the Golgi; Exo1 ^59^ and Exo2 ^60^, which prevent proteins from leaving the endoplasmic reticulum; and tunicamycin, which disrupts normal protein processing in the ER, provides strong and convergent evidence that this pathway plays no role in HSPA1A’s journey to the plasma membrane (Fig. 2 and Supplemental Fig. S2). To block ER-to-Golgi transport, cells were treated with Brefeldin A (BFA), which inhibits ARF1 activation and causes Golgi collapse, or with Exo1 and Exo2, which interfere with early ER export through distinct mechanisms that do not involve ARF1-GEF inhibition. None of these treatments significantly altered HSPA1A PM localization at either 0h or 8h recovery compared to DMSO controls (all p>0.05; Fig. 2). To independently confirm ER-Golgi independence through a mechanistically distinct approach, cells were treated with tunicamycin, an inhibitor of N-linked protein glycosylation that induces ER stress and broadly disrupts conventional secretory cargo processing. Tunicamycin treatment had no significant effect on HSPA1A PM localization at either 0h or 8h recovery compared to controls (Fig. 2). The convergent negative result across four inhibitors targeting mechanistically distinct steps of the conventional secretory pathway, ER export, ARF1-dependent COPI assembly, and glycosylation, makes it unlikely that this negative result reflects incomplete pathway inhibition and collectively establishes that HSPA1A PM localization is independent of the ER-Golgi route.

**Fig. 2.**
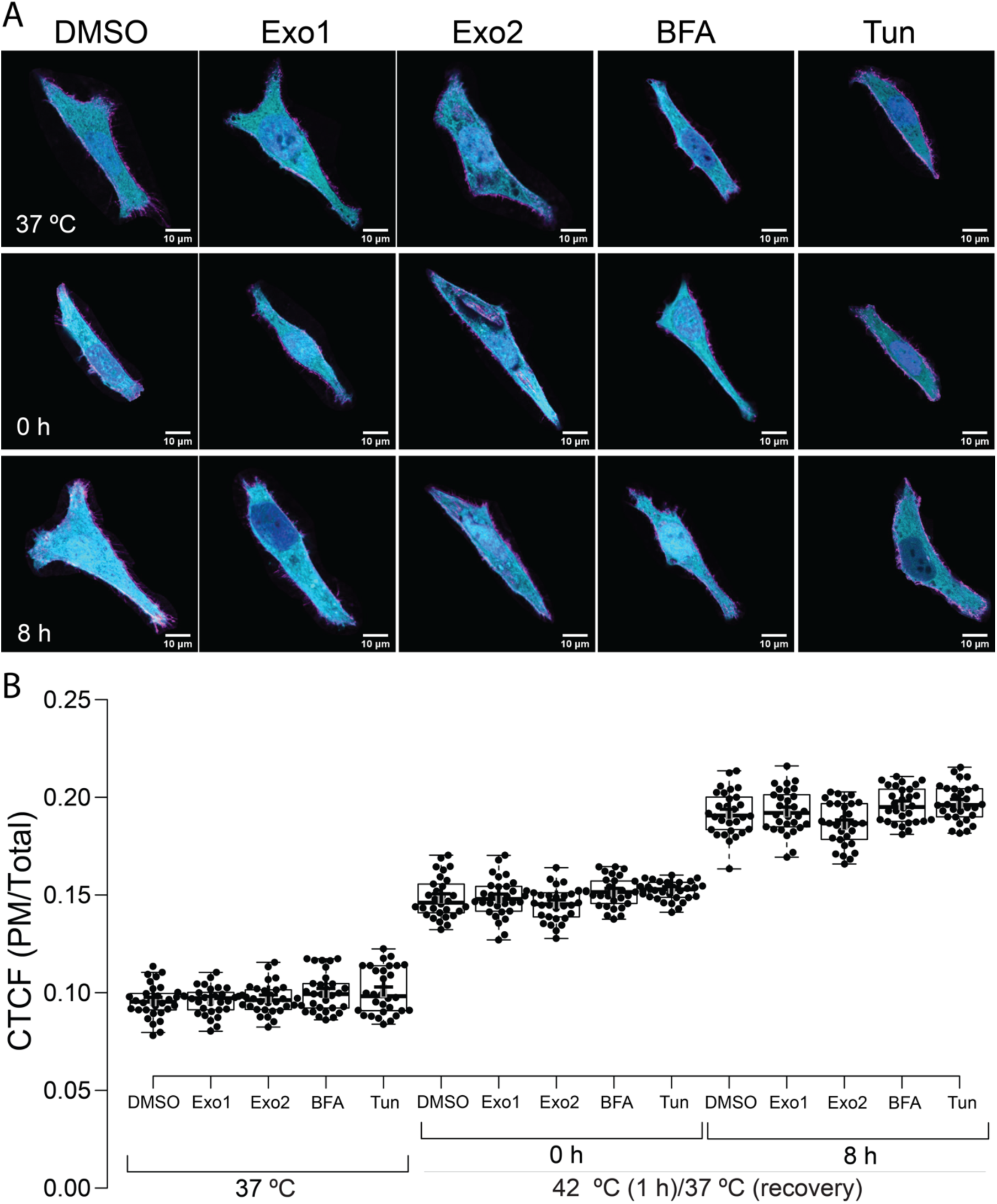
HSPA1A plasma membrane localization is independent of the ER-Golgi secretory pathway. HeLa cells expressing HSPA1A-GFP were treated with DMSO (vehicle control), Exo1, Exo2, Brefeldin A (BFA), or Tunicamycin (Tun) and maintained at 37°C or subjected to heat shock (42°C, 1h) followed by recovery at 37°C for 0h or 8h. **(A)** Representative confocal images show merged channels: HSPA1A-GFP (cyan), plasma membrane marker WGA-FA555 (magenta; FA555 refers to the fluorophore conjugates), and DAPI nuclear stain (blue). Scale bar = 10 μm. Individual channel images for all conditions are shown in Supplemental Fig. S2. **(B)** Quantification of CTCF (PM/Total) for all conditions and timepoints. Each data point represents one cell. Boxes show interquartile range with median; whiskers extend to 1.5× IQR; crosses represent sample means. No statistically significant differences were detected between vehicle control and any drug treatment at any time point (one-way ANOVA with Tukey HSD and Bonferroni post-hoc tests). n=30 cells per condition from three independent experiments.

These findings are consistent with HSPA1A’s lack of canonical secretory signals and place its PM translocation within the broader framework of unconventional protein surface localization and secretion pathways utilized by other leaderless cytosolic proteins ^9,10^.

### 3.3 Endosomal and Lysosomal Pathways Contribute to HSPA1A Plasma Membrane Localization

Having established that HSPA1A does not rely on the ER-Golgi pathway and that it redistributes across endosomal and lysosomal compartments following heat shock, we next asked whether disruption of these pathways functionally affects HSPA1A PM localization. To address this, cells were treated with inhibitors targeting distinct steps of endosomal maturation and lysosomal function, and HSPA1A PM localization was quantified at 0h and 8h of recovery (Fig. 3 and Supplemental Fig. S3).

**Fig. 3.**
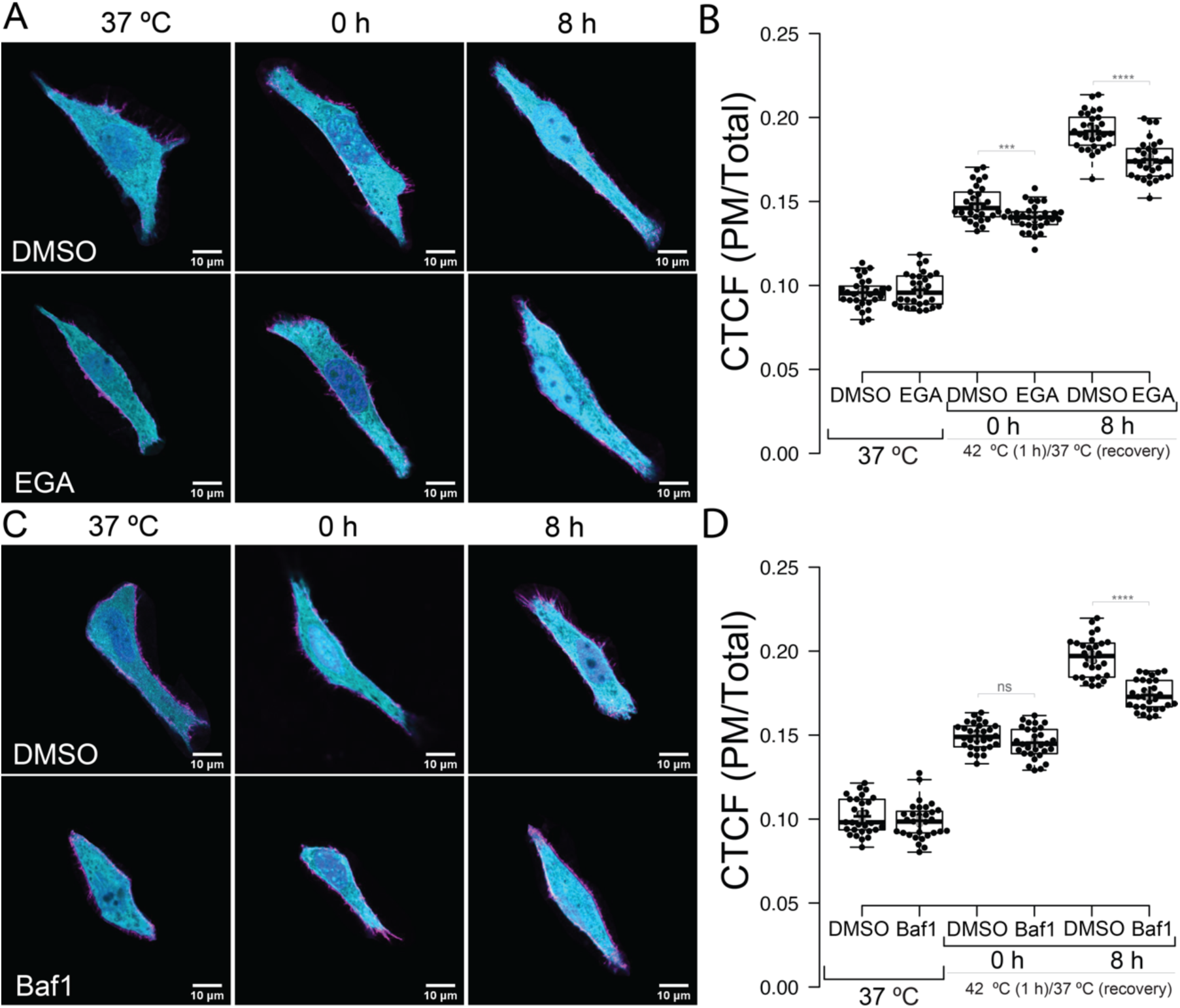
Endosomal maturation and lysosomal function contribute to HSPA1A plasma membrane localization. HeLa cells expressing HSPA1A-GFP were treated with DMSO (vehicle control), EGA, or Bafilomycin A1 (BafA1) and maintained at 37°C or subjected to heat shock (42°C, 1h) followed by recovery at 37°C for 0h or 8h. Representative confocal images show merged channels: HSPA1A-GFP (cyan), WGA-FA555 (magenta), and DAPI (blue). Scale bar = 10 μm. **(A)** Representative images for DMSO and EGA conditions. **(B)** Quantification of CTCF (PM/Total) for EGA and DMSO conditions. **(C)** Representative images for DMSO and BafA1 conditions. **(D)** Quantification of CTCF (PM/Total) for BafA1 and DMSO conditions. Each data point represents one cell. Boxes show interquartile range with median; whiskers extend to 1.5× IQR. Statistical comparisons by one-way ANOVA with Tukey’s post-hoc test. *** p<0.001, **** p<0.0001. Comparisons not indicated were not statistically significant. n=30 cells per condition from three independent experiments.

To test whether progression through the endosomal system is required for HSPA1A PM localization, cells were treated with EGA, a selective inhibitor of early-to-late endosome trafficking that blocks endosomal maturation without impairing endocytosis or lysosomal acidification ^61^. EGA treatment had no significant effect under basal conditions (37°C; p=0.49) but resulted in a small yet statistically significant reduction in HSPA1A PM localization at both 0h recovery (5.5% reduction vs. DMSO control, p=0.0008) and 8h recovery (8.9% reduction, p<0.0001; Fig. 3). This finding supports a contribution of endosomal maturation to HSPA1A PM delivery, while the modest magnitude of the effect suggests that no single endosomal route is solely responsible.

To assess whether lysosomal acidification and degradative activity contribute to HSPA1A PM localization, cells were treated with Bafilomycin A1 (BafA1), which inhibits the vacuolar-type H⁺-ATPase (V-ATPase) ^62^, preventing acidification of endosomes and lysosomes, impairing degradative activity, and blocking autophagosome-lysosome fusion. BafA1 treatment had no significant effect at 37°C (p=0.34) or at 0h recovery (p=0.08), but produced a significant reduction at 8h recovery (11.5% reduction vs. DMSO, p<0.0001; Fig. 3 and Supplemental Fig. S3). The restriction of this effect to the later recovery time point is consistent with the progressive accumulation of HSPA1A in LAMP1-positive compartments observed in section 3.1, suggesting that lysosomal involvement in HSPA1A trafficking becomes more prominent as recovery proceeds.

Subcellular fractionation and western blotting provided qualitative biochemical support for the redistribution of HSPA1A into PM-, lysosome-, and early endosome-enriched fractions following heat shock, with patterns consistent with the colocalization data described above (Supplemental Fig. S4).

The persistence of HSPA1A in lysosomal and endosomal fractions beyond the point of peak PM localization argues against a simple model of complete lysosomal exocytosis. Instead, these findings suggest that HSPA1A trafficking involves bifurcated fates: a fraction is delivered to the PM during recovery, while another fraction is retained within late endosomal and lysosomal compartments, potentially for delayed release, regulated recycling, or degradation. Taken together, the pharmacological and biochemical data support a model in which multiple endosomal and lysosomal routes contribute in parallel to HSPA1A PM localization, with no single pathway acting as a dominant or exclusive route.

### 3.4 Lysosomal Lipid Composition Regulates HSPA1A Plasma Membrane Localization

The progressive accumulation of HSPA1A in LAMP1-positive compartments following heat shock, together with the modest but significant effect of BafA1 on PM localization, raised the question of whether lysosomal compartments actively regulate HSPA1A trafficking rather than simply representing a retention or degradation fate. To address this, we asked whether the lipid composition of late endosomal and lysosomal membranes, specifically the content of bis(monoacylglycerol)phosphate (BMP; a lipid enriched in the internal membranes of late endosomes and multivesicular bodies ^63^), influences HSPA1A PM localization.

To first establish whether BMP levels change dynamically during heat shock recovery, intracellular BMP immunoreactivity was quantified by ICC ^16,55^ using the anti-BMP antibody at 37°C at 0h and 8h recovery. BMP immunoreactivity increased significantly and progressively following heat shock, by 27.7% at 0h recovery (p<0.0001) and 39.2% at 8h recovery (p<0.0001) compared to unstressed cells, with continued accumulation between the two recovery timepoints (9.1%, p<0.0001; Fig. 4A, 4B, and Supplemental Fig. S5). This progressive increase in lysosomal BMP immunoreactivity parallels the accumulation of HSPA1A in LAMP1-positive compartments described in section 3.1 and is consistent with heat shock-induced lipid remodeling creating a permissive lysosomal environment for HSPA1A trafficking.

**Fig. 4.**
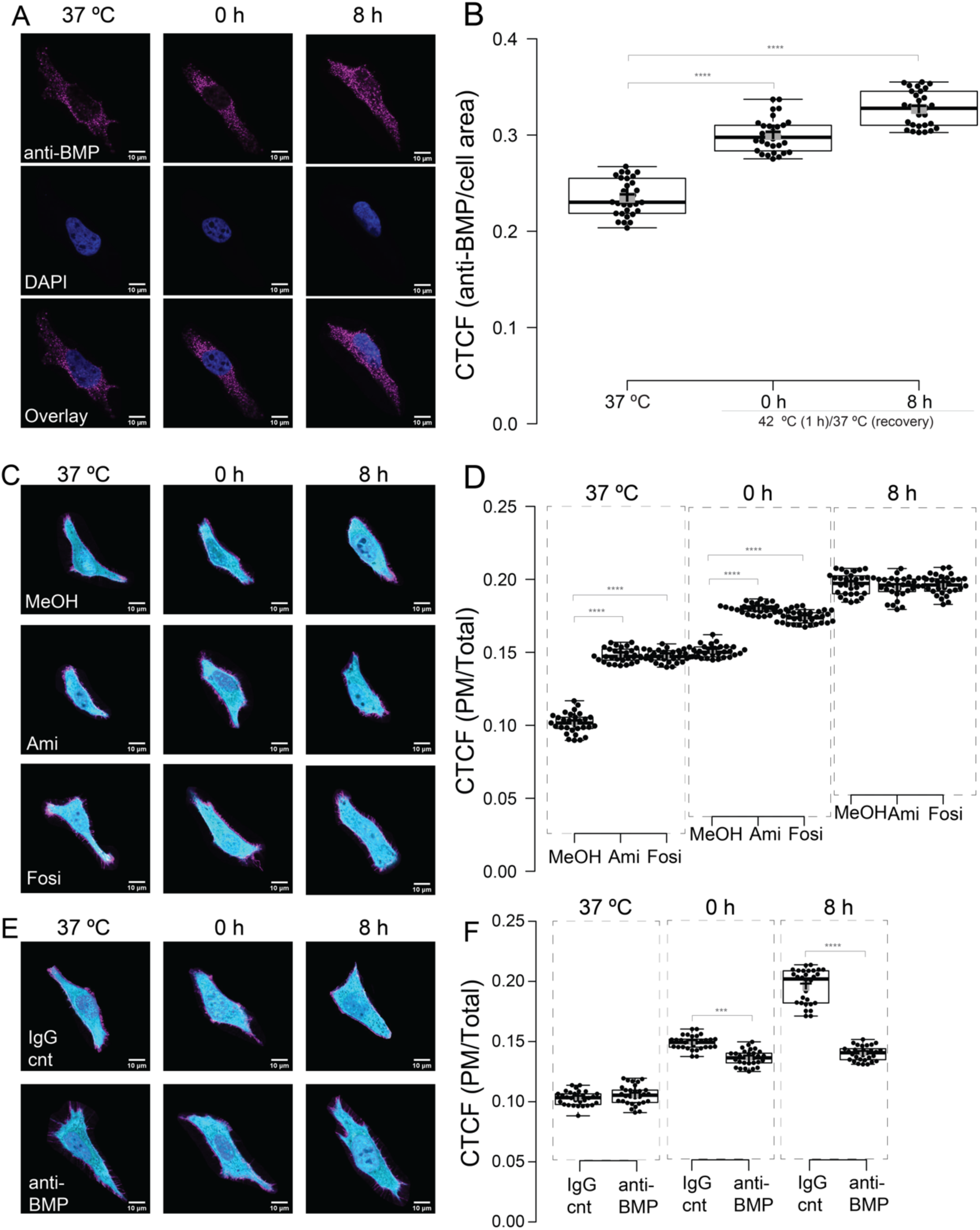
Lysosomal BMP content regulates HSPA1A plasma membrane localization. **(A)** HeLa cells were maintained at 37°C or subjected to heat shock (42°C, 1h) followed by recovery at 37°C for 0h or 8h. Representative ICC images showing intracellular BMP immunoreactivity using anti-BMP antibody (clone 6C4; magenta), and DAPI (blue). Scale bar = 10 μm. **(B)** Quantification of CTCF (anti-BMP/cell area), showing progressive increase in lysosomal BMP immunoreactivity following heat shock. **(C)** HeLa cells expressing HSPA1A-GFP were maintained at 37°C or subjected to heat shock (42°C, 1h) followed by recovery at 37°C for 0h or 8h. Representative confocal images show merged channels: HSPA1A-GFP (cyan), WGA-FA555 (magenta), and DAPI (blue). Scale bar = 10 μm. Representative images of cells treated with methanol vehicle control (MeOH), Amiodarone (Ami), or Fosinopril (Fosi). **(D)** Quantification of CTCF (PM/Total) for MeOH, Ami, and Fosi conditions across all timepoints. **(E)** Representative images of cells delivered a mouse IgG isotype control antibody (IgG-cnt) or anti-BMP antibody via BioPORTER. **(F)** Quantification of CTCF (PM/Total) for IgG-cnt and anti-BMP conditions across all timepoints. Each data point represents one cell. Boxes show interquartile range with median; whiskers extend to 1.5× IQR. Statistical comparisons by one-way ANOVA with Tukey’s post-hoc test. *** p<0.001, **** p<0.0001. Comparisons not indicated were not statistically significant. n=30 cells per condition for all panels from three independent experiments.

To test whether elevated lysosomal BMP levels are sufficient to enhance HSPA1A PM localization, cells were treated with Amiodarone or Fosinopril ^64,65^, two pharmacological agents known to promote BMP accumulation in lysosomal compartments through inhibition of lysosomal phospholipase activity (Fig. 4 and Supplemental Fig. S5). Treatment with either compound significantly increased HSPA1A PM localization under basal conditions (37°C) compared to methanol vehicle controls, Amiodarone by 46.0% (p<0.0001) and Fosinopril by 45.4% (p<0.0001), demonstrating that BMP enrichment alone is sufficient to drive HSPA1A to the PM even in the complete absence of heat shock (Fig. 4). At 0h post-heat shock, both compounds produced further significant increases compared to vehicle controls (Amiodarone: +19.6%, p<0.0001; Fosinopril: +15.2%, p<0.0001). At 8h recovery, no significant difference was observed between drug-treated and vehicle control cells (Amiodarone: p=0.29; Fosinopril: p=0.77), consistent with a ceiling effect at this timepoint where heat shock-induced PM localization is already near maximal (Fig. 4).

To determine whether BMP is not merely correlative but functionally necessary for HSPA1A PM localization, an anti-BMP antibody was delivered intracellularly using BioPORTER reagent. A mouse IgG isotype control antibody, delivered under identical conditions, served as a control to account for the nonspecific effects of the delivery reagent and the presence of intracellular antibody. The specificity of the BioPORTER delivery approach was further confirmed by delivering an anti-PS antibody under identical conditions, which completely abolished heat shock-induced HSPA1A PM localization (Supplemental Fig. S6), consistent with the previously established essential role of PS ^25^. Anti-BMP delivery had no significant effect under basal conditions (37°C; p=0.17), consistent with the low baseline PM localization of HSPA1A in unstressed cells. Following heat shock, anti-BMP delivery produced a significant reduction in HSPA1A PM localization at 0h recovery (8.6% reduction vs. isotype control, p<0.0001) and a more substantial reduction at 8h recovery (28.5% reduction, p<0.0001; Fig. 4).

The gain-of-function result (BMP enrichment driving a ∼46% increase in PM localization at 37°C) combined with the loss-of-function result (BMP blockade reducing PM localization by ∼29% at 8h) establishes BMP as a functionally required regulator of HSPA1A PM localization rather than a passive correlate of lysosomal identity. Notably, the ability of BMP enrichment to drive HSPA1A PM localization independently of heat shock indicates that lipid permissiveness, specifically the BMP content of lysosomal compartments, can substitute for the stress trigger under appropriate conditions. The distinction between the strong effect of BMP modulation and the modest effect of BafA1 indicates that lysosomal lipid composition, rather than lysosomal acidification or degradative activity, is the primary lysosomal regulatory input into HSPA1A trafficking. This positions BMP-enriched lysosomal compartments not as terminal degradative destinations but as active regulatory hubs whose lipid state determines their competence for PM localization. The regulatory role of BMP at the lysosomal level raises the broader question of how lipid identity is maintained and coordinated across the entire endosomal network through which HSPA1A traffics. This question was directly addressed by the compartment-specific PI(4)P depletion experiments presented below.

### 3.5 PI(4)P Is Required Across Multiple Endosomal Compartments for HSPA1A Plasma Membrane Localization

Previous work from our laboratory established that PI(4)P at the plasma membrane is essential for HSPA1A PM localization, and that heat shock drives PI(4)P accumulation at the PM through activation of PI4KIIIα ^26,27^. Those studies used a PM-anchored Sac1 phosphatase system and pharmacological PI(4)P depletion to demonstrate PI(4)P dependency at the cell surface. However, whether PI(4)P within endosomal compartments, through which HSPA1A traffics prior to PM delivery, is also required remains unknown. To address this, we used a rapamycin-inducible dimerization system to recruit the Sac1 phosphatase specifically to Rab5-positive early endosomes, Rab7-positive late endosomes, and lysosomal compartments, and assessed the effect on HSPA1A PM localization following heat shock.

To test whether PI(4)P within early endosomal compartments contributes to HSPA1A trafficking, Sac1-WT was recruited to Rab5-positive endosomes by rapamycin-induced dimerization of FKBP-tagged Sac1 with FRB-tagged Rab5. Recruitment of active Sac1-WT to Rab5-positive compartments resulted in a significant reduction in HSPA1A PM localization at both 0h recovery (10.1% reduction vs. Sac1-Dead+Rapa control, p<0.0001) and 8h recovery (13.5% reduction, p<0.0001; Fig. 5 and Supplemental Fig. S6). Control conditions included Sac1-WT without rapamycin, confirming that Sac1 expression alone without compartment recruitment has no effect (0h: p=0.35; 8h: p=0.998), and the catalytically inactive Sac1-Dead construct with and without rapamycin, confirming that the reduction in PM localization requires phosphatase activity rather than simply reflecting overexpression or steric effects of the recruited protein. No significant difference was observed between Sac1-WT and Sac1-Dead under basal conditions (37°C: p=0.61), confirming that the effect is heat shock-dependent. These results demonstrate that PI(4)P within early endosomes is functionally required for efficient HSPA1A trafficking to the PM, consistent with PI(4)P’s role in defining endosomal membrane identity and supporting vesicular cargo sorting ^66,67^.

**Fig. 5.**
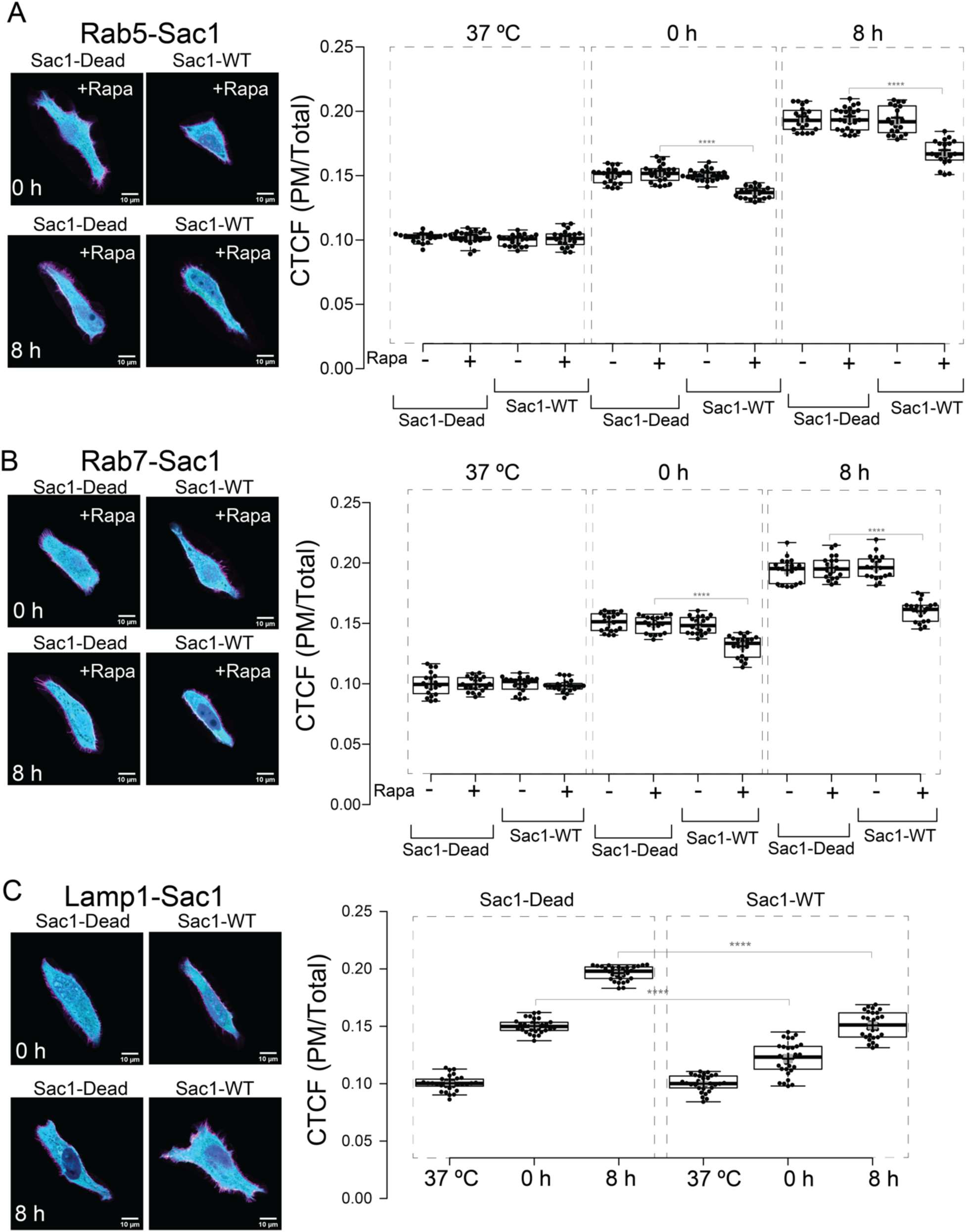
PI(4)P is required across multiple endosomal compartments for HSPA1A plasma membrane localization. HeLa cells co-expressing HSPA1A-GFP with rapamycin-inducible FRB-tagged compartment anchors and FKBP-tagged Sac1 phosphatase (wild-type, Sac1-WT; or catalytically inactive, Sac1-Dead) were maintained at 37°C or subjected to heat shock (42°C, 1h) followed by recovery at 37°C for 0h or 8h. For Rab5 and Rab7 experiments, rapamycin (+Rapa) was added where indicated to induce dimerization and recruit Sac1 to the target compartment. Controls included Sac1-WT without rapamycin and Sac1-Dead with rapamycin, confirming that Sac1 expression alone and compartment recruitment, in the absence of phosphatase activity, do not affect HSPA1A PM localization. For lysosomal experiments, constitutively lysosome-targeted Sac1-WT and Sac1-Dead constructs were used in the absence of rapamycin. Representative confocal images show merged channels: HSPA1A-GFP (cyan), WGA-FA555 plasma membrane marker (magenta), and DAPI nuclear stain (blue). Scale bar = 10 μm. **(A)** Recruitment of Sac1 to Rab5-positive early endosomes: representative images (left) and quantification of CTCF (PM/Total) (right). **(B)** Recruitment of Sac1 to Rab7-positive late endosomes: representative images (left) and quantification of CTCF (PM/Total) (right). **(C)** Recruitment of Sac1 to lysosomal compartments using lysosome-targeted Sac1-WT and Sac1-Dead constructs: representative images (left) and quantification of CTCF (PM/Total) (right). Each data point represents one cell. Boxes show interquartile range with median; whiskers extend to 1.5× IQR. Statistical comparisons by one-way ANOVA with Tukey’s post-hoc test. **** p<0.0001. Comparisons not indicated were not statistically significant. n=30 cells per condition from three independent experiments. Individual channel images for all conditions are shown in Supplemental Fig. S7.

**Fig. 6.**
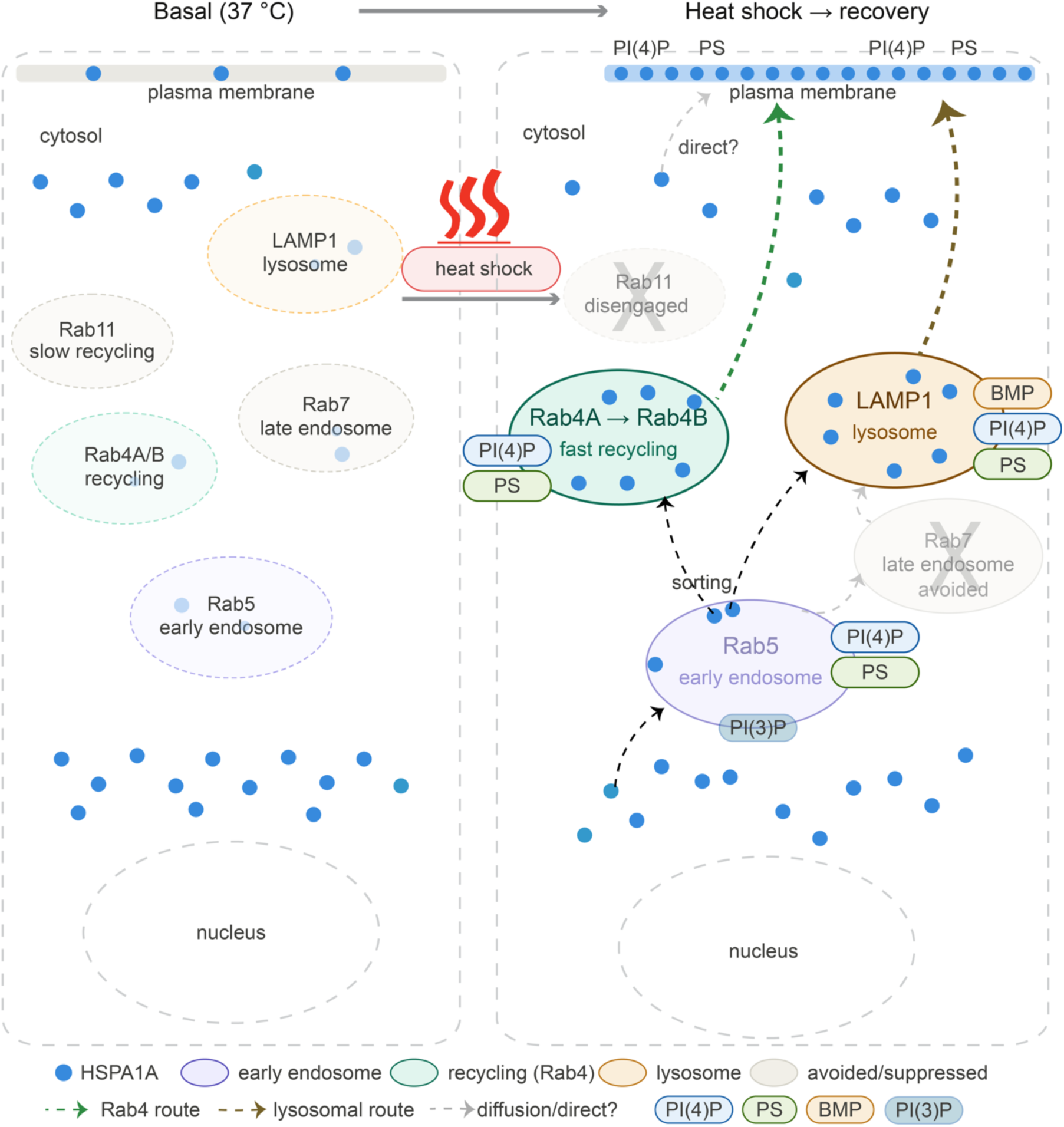
Proposed model of lipid-gated vesicular trafficking of HSPA1A to the plasma membrane. Schematic comparing HSPA1A distribution and trafficking routes under basal conditions (37 °C, left) and following heat shock (42 °C, 1h) with recovery (right). Under basal conditions, HSPA1A resides predominantly in the cytosol with minimal plasma membrane (PM) localization; endosomal and lysosomal compartments are present but not enriched for HSPA1A. Following heat shock, stress-induced lipid remodeling drives recruitment of cytosolic HSPA1A to PI(3)P-enriched Rab5-positive early endosomes, which serve as the initial sorting station. From Rab5, HSPA1A is directed along two parallel forward routes: a fast-recycling route through Rab4A- then Rab4B-positive endosomes, and a lysosomal route through BMP-enriched LAMP1-positive compartments. Both routes converge at the plasma membrane, where PI(4)P and phosphatidylserine (PS) are required for final docking and stable membrane association. PI(4)P is required not only at the PM but at multiple endosomal stages, including Rab5-positive early endosomes and lysosomal compartments (Fig. 5), establishing a distributed lipid requirement throughout the trafficking route. PS is present at endosomal membranes and is essential at the PM docking step. BMP enrichment at lysosomal compartments regulates PM localization competence (Fig. 4). Rab7-positive late endosomes and Rab11-positive slow recycling endosomes are actively avoided and disengaged, respectively, following heat shock. A minor non-vesicular contribution via direct diffusion and lipid-dependent PM capture is also possible but not directly tested (dashed gray arrow). The classical ER-Golgi secretory pathway is not required (Fig. 2). Blue circles, HSPA1A; compartment colors as indicated in legend.

To determine whether the PI(4)P requirement extends beyond early endosomes to late endosomal compartments, the same rapamycin-inducible system was used to recruit Sac1-WT to Rab7-positive late endosomes. Recruitment of active Sac1-WT to Rab7-positive compartments resulted in a significant reduction in HSPA1A PM localization at both 0h recovery (12.2% reduction vs. Sac1-Dead+Rapa control, p<0.0001) and 8h recovery (18.3% reduction, p<0.0001; Fig. 5). The same specificity controls, Sac1-WT without rapamycin and Sac1-Dead with and without rapamycin, confirmed that the effect is dependent on both compartment recruitment and catalytic activity (Dead +Rapa vs. -Rapa: 0h p=0.42, 8h p=0.53; 37°C baseline: p=0.54). These results demonstrate that PI(4)P within late endosomes is independently required for HSPA1A PM localization, extending the PI(4)P requirement beyond the early endosomal sorting step to a later stage of the trafficking route.

To assess whether the PI(4)P requirement extends further to lysosomal compartments, Sac1 was recruited to lysosomal membranes using a lysosome-targeted FRB construct. Lysosomal Sac1-WT recruitment resulted in a statistically significant reduction in HSPA1A PM localization at both 0h recovery (18.9% reduction vs. Dead mutant control, p<0.0001) and 8h recovery (23.2% reduction, p<0.0001; Fig. 5 and Supplemental Fig. S6). No significant difference was observed between wild type and dead mutant Sac1 under basal conditions (37°C: p=0.87), confirming heat shock-dependence and specificity of the effect. The effect magnitude at the lysosomal compartment was comparable to that observed at Rab7 and larger than at Rab5, consistent with the progressive importance of PI(4)P at successive stages of the trafficking route and the parallel contribution of BMP as the dominant lipid regulator at the lysosomal level. Catalytically inactive Sac1-Dead controls confirmed that the reduction requires phosphatase activity rather than simply reflecting overexpression or compartment targeting artifacts.

The comparable reductions in HSPA1A PM localization following Sac1 recruitment to Rab5-positive, Rab7-positive, and lysosomal compartments (each independently significant, each dependent on catalytic activity, and together revealing a gradient of effect sizes from ∼10% at early endosomes to ∼23% at lysosomes) collectively argue against a model in which PI(4)P functions solely as a terminal docking cue at the plasma membrane. Instead, these data support a model in which PI(4)P establishes a permissive membrane state at multiple stages of the endosomal network, contributing to HSPA1A sorting competence at early endosomes, to trafficking progression through late endosomes, and to regulation of PM localization at lysosomal compartments. This gradient is consistent with PI(4)P becoming progressively more important as HSPA1A approaches the PM, with the dominant requirement at the final docking step as established previously ^26,27^. This distributed PI(4)P requirement is consistent with the known role of phosphoinositides in defining membrane identity and coordinating vesicular trafficking specificity across compartments ^37,66,67^.

Taken together with the published findings that PI(4)P at the PM is essential for final HSPA1A docking ^26,27^, the present results establish PI(4)P as a lipid signal required continuously throughout the HSPA1A trafficking route, from entry into early endosomes through late endosomal and lysosomal intermediates to final plasma membrane association. These findings represent a significant extension of the current model of lipid-dependent HSPA1A trafficking and identify compartment-specific PI(4)P regulation as a novel layer of control over this unconventional secretory pathway.

## 4. Discussion

HSPA1A is a leaderless molecular chaperone that localizes to the plasma membrane of heat-shocked and cancer cells, where it contributes to therapeutic resistance, membrane stabilization, and immune modulation ^6,21,68–73^. Despite its clinical significance, the mechanisms underlying its PM translocation have remained poorly defined. In this study, we investigated the vesicular route and lipid determinants of HSPA1A trafficking to the PM following heat shock. Our findings support a model in which HSPA1A is trafficked through the endo-lysosomal network via a lipid-dependent vesicular mechanism that bypasses the classical ER-Golgi secretory pathway. This mechanism involves coordinated redistribution across multiple endosomal compartments and is regulated at the lysosomal level by BMP-dependent PM competence and at multiple endosomal stages by PI(4)P-dependent membrane identity. Together with previously published findings establishing PS and PI(4)P as essential PM docking lipids ^5,8,18,25–27,43,74^, these results define a lipid-gated vesicular trafficking framework for HSPA1A PM localization and extend the current model of unconventional chaperone secretion.

Confocal imaging and colocalization analysis revealed that heat shock triggers a coordinated and selective redistribution of HSPA1A across intracellular compartments. The temporal pattern is inconsistent with passive diffusion or nonspecific membrane association. Instead, it suggests active sorting into specific vesicular pathways in a temporally regulated manner. HSPA1A transiently associates with PI(3)P-enriched FYVE-positive endosomes early in the response, rapidly engages Rab4A-positive fast recycling endosomes, shifts toward Rab4B-positive compartments at later timepoints, and progressively accumulates in LAMP1-positive lysosomes. The concurrent decrease in Rab5, Rab7, and Rab11 association indicates that HSPA1A does not follow canonical early endosomal retention, degradative late endosomal, or slow recycling routes following stress. The temporal distinction between Rab4A and Rab4B involvement is particularly noteworthy. Rab4A is associated with immediate fast recycling from early endosomes, while Rab4B has been linked to sustained recycling during later recovery phases ^75^. The sequential engagement of these two GTPases is consistent with a progressive forward trafficking model in which HSPA1A is directed through recycling compartments toward the PM over the course of the recovery period. This pattern of compartmental redistribution is consistent with Type II or Type III unconventional protein secretion, in which leaderless cytosolic proteins engage the endo-lysosomal network to reach the PM or extracellular space ^9,10^. The absence of increased FYVE-HSPA1A colocalization at 0h, despite the established functional requirement for PI(3)P in HSPA1A PM localization ^27^, is consistent with competition between the FYVE domain and HSPA1A for the same PI(3)P-enriched membrane sites. This interpretation is directly supported by the approximately 50% loss of PM localization observed when PI(3)P is masked by EEA1 overexpression or depleted by wortmannin ^27^. This competition model reconciles the functional PI(3)P requirement with the colocalization dynamics and reinforces the notion that HSPA1A’s association with PI(3)P endosomes is transient and rapidly resolved as the protein progresses through the trafficking network. The selective engagement of these compartments and the avoidance of canonical degradative and slow-recycling routes are consistent with a directed, unconventional trafficking mechanism rather than passive membrane association. This conclusion is directly supported by the insensitivity of HSPA1A PM localization to ER-Golgi disruption.

The absence of any effect on HSPA1A PM localization across four inhibitors that each block a different step of the classical secretory pathway provides strong and convergent evidence that this pathway plays no role in HSPA1A’s journey to the plasma membrane. BFA collapses the Golgi. Exo1 and Exo2 prevent proteins from leaving the endoplasmic reticulum. Tunicamycin disrupts normal ER protein processing. None produced a significant effect at any of the time points tested. These findings are consistent with HSPA1A’s lack of a signal peptide and align with prior observations that HSPA1A PM localization is insensitive to disruption of the secretory pathway ^11–17^. Together, these results place HSPA1A firmly within the unconventional secretion framework and direct attention toward endosomal and lysosomal routes as the relevant trafficking intermediates. These findings also extend earlier studies examining HSPA1A unconventional secretion. Previous investigations reported that extracellular HSPA1A release is largely insensitive to disruption of the classical secretory pathway, supporting its classification as a leaderless protein exported via non-canonical mechanisms. However, those studies primarily measured HSPA1A in the extracellular medium and did not address the route by which HSPA1A becomes associated with the plasma membrane. Our results demonstrate that the PM-associated pool is likewise independent of ER-Golgi trafficking, while further identifying specific endosomal compartments, lysosomal lipid regulators, and compartmentalized phosphoinositide requirements that govern HSPA1A localization at the cell surface. With the conventional secretory pathway excluded, attention turns to the specific endosomal and lysosomal compartments that actively regulate PM-HSPA1A, among which lysosomal lipid composition emerges as a critical determinant.

Among the lipids that regulate vesicular trafficking within the endo-lysosomal system, bis(monoacylglycerol)phosphate (BMP) stands out as a lipid uniquely enriched in the internal membranes of lysosomes, where it plays structural and regulatory roles in vesicle maturation and cargo sorting. Our results identify it as a key regulator of HSPA1A PM localization. Amiodarone and Fosinopril drive HSPA1A PM localization at 37°C in the complete absence of heat shock, demonstrating that BMP enrichment alone is sufficient to bypass the stress requirement and promote HSPA1A PM localization. Conversely, intracellular blockade of BMP with a functionally delivered anti-BMP antibody significantly reduced HSPA1A PM localization at both 0h and 8h recovery following heat shock, with the effect being more pronounced at 8 h. The functional hierarchy between lipids is also informative. Intracellular delivery of anti-PS antibody via BioPORTER completely abolished heat shock-induced HSPA1A PM localization; HSPA1A remained at unstressed baseline levels across all timepoints, while anti-BMP delivery produced a significant but partial reduction. This distinction is consistent with PS acting as an essential PM docking lipid whose blockade prevents any surface accumulation regardless of upstream trafficking. At the same time, BMP governs lysosomal PM localization competence in a modulatory capacity, with partial redundancy or parallel routes accounting for residual PM localization ^25^.

The convergence of gain-of-function and loss-of-function results identifies BMP as a functionally required regulator rather than a passive correlate of lysosomal identity. Its role here as a trafficking enabler is consistent with BMP’s broader function in regulating vesicle maturation and membrane remodeling within the late endosomal system ^76^. The distinction between the strong effect of BMP modulation and the modest effect of BafA1 is important. It indicates that lysosomal lipid composition, rather than lysosomal acidification or degradative activity, is the primary lysosomal regulatory input into HSPA1A trafficking. This positions BMP-enriched lysosomal compartments not as terminal degradative destinations but as active regulatory hubs whose lipid state determines their competence for PM localization. The stress bypass phenomenon is also noteworthy. BMP enrichment drives PM localization in the absence of heat shock, suggesting that lipid permissiveness and stress signaling converge on a common trafficking pathway and that, under appropriate lipid conditions, the pathway can operate constitutively. This may be relevant to cancer cells, where lysosomal lipid composition is frequently altered, and constitutive mHSPA1A expression is observed ^5,21^. Notably, heat shock itself drives a progressive increase in intracellular BMP immunoreactivity, rising 27.7% by 0h and 39.2% by 8h recovery, a temporal pattern that is distinct from the dynamics of PS and PI(4)P, both of which peak immediately after heat shock and partially decline during recovery ^25,26^. The sustained and progressive increase in BMP parallels the continued accumulation of HSPA1A in LAMP1-positive compartments through the recovery period, consistent with BMP acting as a lysosomal trafficking competence signal that becomes increasingly permissive as recovery proceeds rather than a transient stress signal. The regulatory role of BMP at the lysosomal level raises the broader question of how lipid identity is maintained and coordinated across the entire endosomal network through which HSPA1A traffics. This question is directly addressed by the compartment-specific PI(4)P depletion experiments.

To determine where along the trafficking pathway PI(4)P is required for HSPA1A PM localization, we used a molecular tool that selectively removes PI(4)P from a specific intracellular compartment at a time, while leaving all other compartments intact. Prior work from our laboratory demonstrated that PI(4)P at the PM is essential for HSPA1A surface localization and that heat shock drives PI(4)P accumulation through PI4KIIIα activation ^26,27^. Those studies used a PM-anchored phosphatase system and therefore addressed only the terminal docking step. The present findings reveal that PI(4)P depletion within Rab5-positive early endosomes, Rab7-positive late endosomes, and lysosomal compartments each independently reduces HSPA1A PM localization. Effects were dependent on phosphatase catalytic activity, as confirmed by Sac1-Dead controls in all three experiments. Rather than uniform reductions across compartments, the data reveal a gradient of effect sizes. The reduction is smallest at early endosomes, intermediate at late endosomes, and largest at lysosomes. Together, these results argue against a strictly linear model in which PI(4)P functions only at one defined trafficking step. Instead, they support a distributed lipid requirement in which PI(4)P establishes a permissive membrane state at multiple stages of the endosomal network, becoming progressively more important as HSPA1A approaches the plasma membrane. This is consistent with the established role of phosphoinositides in defining compartment identity and coordinating the recruitment of trafficking effectors throughout the vesicular system ^37,66,67^. Recent studies further support the existence of functionally distinct PI(4)P pools within endosomal and lysosomal compartments, consistent with the compartment-specific effects observed here ^40–42^. The substantial but incomplete effect of lysosomal Sac1 recruitment, may reflect the parallel contribution of BMP at this stage, partial redundancy between routes, or differences in PI(4)P pool accessibility. These possibilities are not mutually exclusive and warrant further investigation. Taken together, the Sac1 data reframe PI(4)P not merely as a PM docking cue but as a lipid signal required continuously throughout the HSPA1A trafficking route, from early endosomal sorting through late endosomal and lysosomal intermediates to final PM association. The endosomal redistribution data, ER-Golgi independence, BMP regulatory findings, and distributed PI(4)P requirement can be integrated into a unified model of lipid-gated vesicular trafficking.

While our data strongly support a lipid-dependent vesicular mechanism for HSPA1A translocation to the plasma membrane, we acknowledge that additional mechanisms may contribute in parallel. Heat shock-induced cytoskeletal remodeling may facilitate the directed movement, positioning, or PM fusion of HSPA1A-containing vesicles, given the well-established role of microtubule- and actin-based motors in endo-lysosomal transport. Additionally, a fraction of cytosolic HSPA1A may reach the PM through stochastic diffusion and preferential capture by lipid-enriched domains, particularly under conditions of elevated PI(4)P and PS following heat shock. These possibilities are not mutually exclusive with the vesicular route described here. Rather, they likely represent parallel or complementary contributions that converge on the same lipid-dependent retention mechanism at the cell surface. Distinguishing the relative contributions of active vesicular delivery, cytoskeletal facilitation, and diffusion-based capture will require live-cell tracking approaches and cytoskeletal perturbation experiments beyond the scope of the present study.

Integrating the findings of this study with published lipid-dependency data, we propose the following model. Following heat shock, stress-induced lipid remodeling, including rapid increases in PS ^25^ and PI(4)P driven by PI4KIIIα activation ^26^, promotes recruitment of cytosolic HSPA1A to intracellular membranes. HSPA1A enters PI(3)P-enriched early endosomes, which serve as an initial sorting or transit station. This is consistent with the transient FYVE enrichment and the partial reduction in PM localization upon PI(3)P depletion reported previously ^27^. From early endosomes, HSPA1A progresses through Rab4A- then Rab4B-positive fast recycling endosomes toward the PM, while a parallel fraction is directed toward BMP-enriched lysosomal compartments. BMP governs the PM localization competence of these lysosomal hubs, determining whether vesicles fuse with the PM or retain their cargo for delayed release or degradation. PI(4)P is required at each of these stages to maintain trafficking competence, likely through its role in membrane identity and effector recruitment. Final PM docking requires PI(4)P and PS as essential anchoring lipids at the inner leaflet of the plasma membrane. The overall process does not rely on the ER-Golgi pathway and involves multiple parallel routes rather than a single dominant trafficking sequence, consistent with the modest but convergent effects observed with individual pathway inhibitors. This model places HSPA1A within a broader framework of lipid-gated unconventional secretion pathways and identifies lysosomal BMP content and distributed endosomal PI(4)P as novel regulatory layers controlling chaperone trafficking during stress.

While multiple orthogonal lines of evidence support this model, several important questions remain. The present study establishes that specific lipids are required at multiple endosomal stages for HSPA1A trafficking, but does not resolve the temporal relationships between lipid remodeling events and HSPA1A recruitment, the quantitative lipid thresholds required for PM localization, or whether PS and PI(4)P act independently or cooperatively at defined steps. Determining the precise temporal order in which PS increases, PI(4)P accumulates, and HSPA1A is recruited to each compartment will be essential for understanding how the trafficking pathway is regulated. Whether these relationships differ between non-transformed and cancer cell backgrounds is an equally important question. The molecular basis of HSPA1A membrane association at each compartment also remains unclear, given the absence of canonical lipid-binding domains. Identifying which regions of HSPA1A mediate interactions with PI(4)P, BMP, and PS and whether these interactions involve specific conformational states, dimerization, or post-translational modifications will be critical for establishing a direct structural link between lipid composition and vesicular entry. Whether PM-localized HSPA1A adopts a defined membrane topology with specific domains exposed extracellularly, and whether this orientation is required for its membrane-protective and immunomodulatory functions, also remains to be determined. Addressing these questions will establish the structural and functional basis of lipid-regulated HSPA1A membrane localization and identify precise molecular targets for disrupting mHSPA1A in cancer cells, where constitutive lysosomal lipid remodeling may sustain its PM-associated pro-survival and therapy-resistance functions.

## 5. Conclusion

This study demonstrates that HSPA1A trafficking to the plasma membrane following heat shock is mediated by lipid-dependent vesicular pathways that bypass the classical ER-Golgi secretory route and engage the endo-lysosomal network in a coordinated, compartment-specific manner. PI(4)P is required not only at the plasma membrane for final docking but across multiple endosomal stages, establishing a distributed lipid requirement that reframes PI(4)P as a continuous trafficking competence signal rather than a terminal anchoring cue. BMP-enriched lysosomal compartments serve as regulatory hubs whose lipid state governs the PM localization of HSPA1A, and together these findings define a lipid-gated unconventional secretion mechanism with implications for cancer cells in which constitutive lysosomal lipid alterations may sustain plasma membrane HSPA1A and its associated pro-survival and therapy-resistance functions.

## Author contributions

**Jensen Low**: Investigation, Data curation, Formal analysis, Visualization, Writing (original draft). **Azalea Blythe Cuaresma**: Investigation, Data curation, Formal analysis, Visualization. **Clarisse K. Martin**: Investigation, Formal analysis, Methodology. **Allen Badolian**: Investigation, Formal analysis. **Maha AlSebaye**: Formal analysis. **Robert V. Stahelin**: Conceptualization, Supervision, Writing, review & editing. **Nikolas Nikolaidis**: Conceptualization, Investigation, Data curation, Methodology, Supervision, Project administration, Funding acquisition, Writing (original draft), Writing (review & editing).

## Abbreviations

ANOVA: Analysis of Variance
BFA: Brefeldin A
BMP: Bis(monoacylglycerol)phosphate
cDNA: Complementary DNA
CTCF: Corrected Total Cell Fluorescence
DMSO: Dimethyl Sulfoxide
FBS: Fetal Bovine Serum
GFP: Green Fluorescent Protein
HSD: Honestly Significant Difference
HSPA1A: Heat Shock Protein A1A
ICC: Immunocytochemistry
IQR: Interquartile Range
MEM: Modified Eagle Medium
NGS: Normal Goat Serum
PBS: Phosphate-Buffered Saline
PFA: Paraformaldehyde
PI(3)P: Phosphatidylinositol 3-Phosphate
PI(4)P: Phosphatidylinositol 4-Phosphate
PM: Plasma Membrane
PS: Phosphatidylserine
RFP: Red Fluorescent Protein
ROI: Region of Interest
RT: Room Temperature
SDS-PAGE: Sodium Dodecyl Sulfate-Polyacrylamide Gel Electrophoresis
UPS: Unconventional Protein Secretion
WGA: Wheat Germ Agglutinin

