## Supplemental Figures S1-S7 for "Lipid-Gated Vesicular Trafficking Directs HSPA1A to the Plasma Membrane Through the Endo-Lysosomal Network"

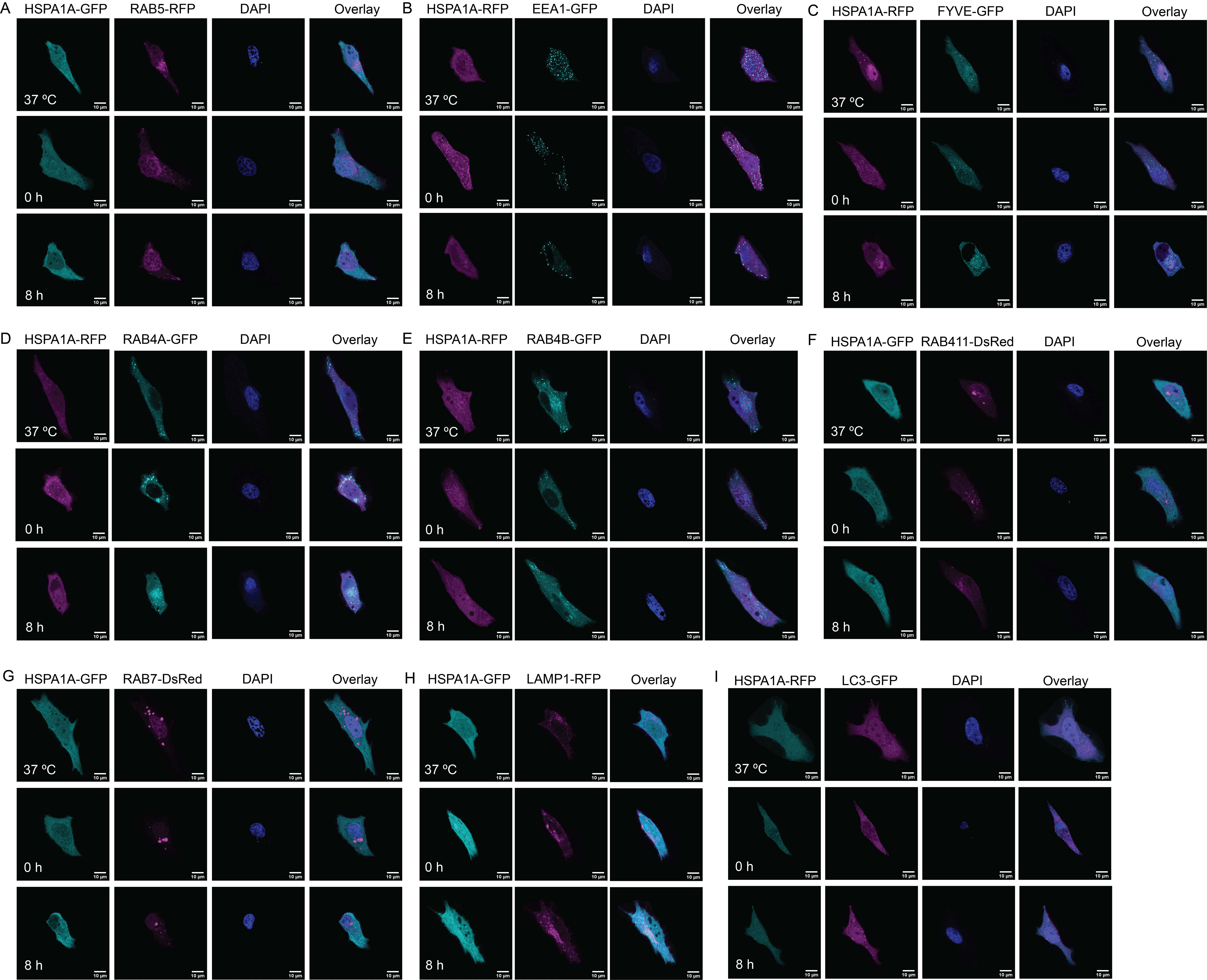
A
HSPA1A-GFP
RAB5-RFP
DAPI
Overlay
B
HSPA1A-RFP
EEA1-GFP
DAPI
Overlay
C
HSPA1A-RFP
FYVE-GFP
DAPI
Overlay
D
HSPA1A-RFP
RAB4A-GFP
DAPI
Overlay
E
HSPA1A-RFP
RAB4B-GFP
DAPI
Overlay
F
HSPA1A-GFP
RAB411-DsRed
DAPI
Overlay
G
HSPA1A-GFP
RAB7-DsRed
DAPI
Overlay
H
HSPA1A-GFP
LAMP1-RFP
Overlay
I
HSPA1A-RFP
LC3-GFP
DAPI
Overlay
37 ºC
0 h
8 h

**Supplemental Fig. S1. Representative confocal images corresponding to Figure 1.** HeLa cells co-expressing HSPA1A-GFP or HSPA1A-RFP with fluorescently tagged compartment markers were maintained at 37°C or subjected to heat shock (42°C, 1h) followed by recovery at 37°C for 0h or 8h. For each marker, individual channels and merged overlay are shown at all three timepoints. Panels correspond to markers shown in Fig. 1: **(A)** RAB5, **(B)** EEA1, **(C)** FYVE, **(D)** RAB4A, **(E)** RAB4B, **(F)** RAB11, **(G)** RAB7, **(H)** LAMP1, **(I)** LC3. Scale bar = 10 μm.

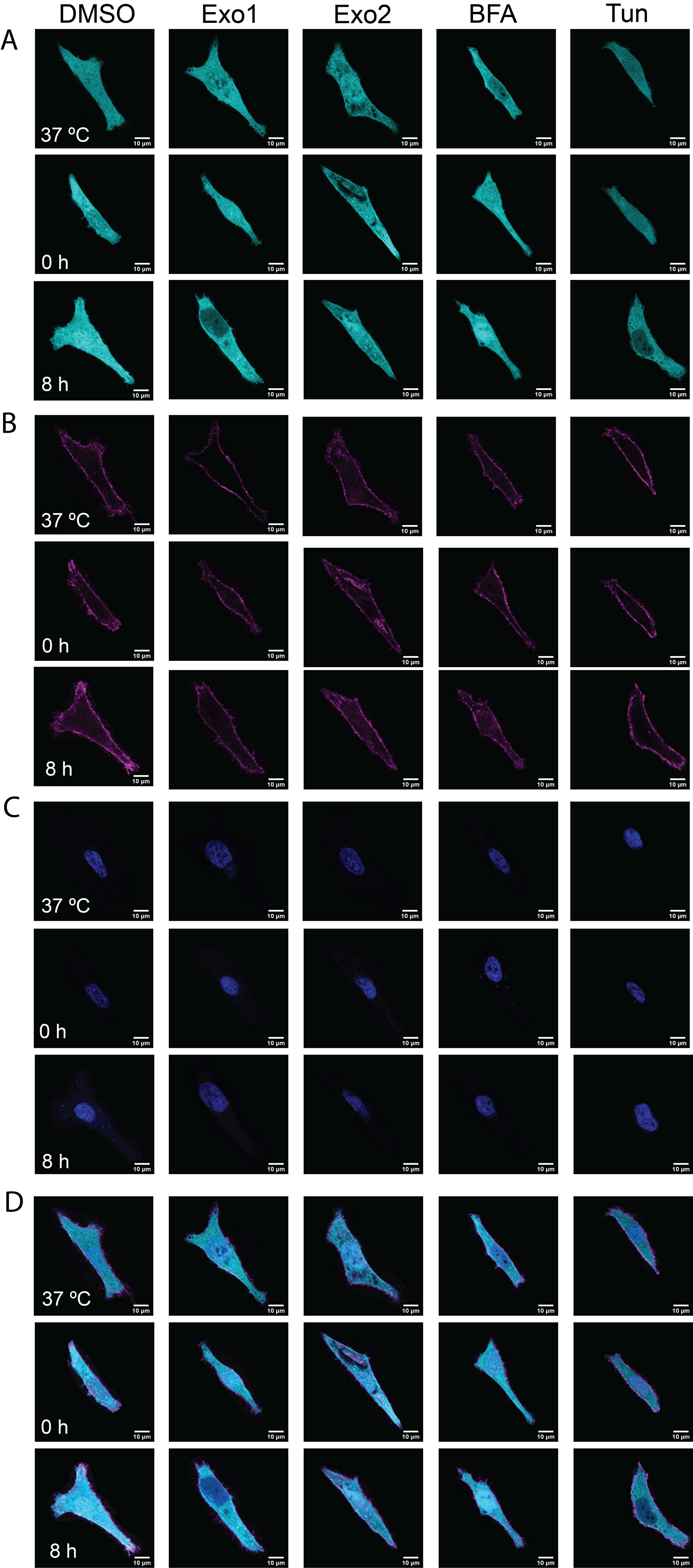
DMSO
Exo1
Exo2
BFA
Tun
A
B
C
D
37 ºC
0 h
8 h
10 µm

**Supplemental Fig. S2. Individual channel images corresponding to Figure 2A. (A)** HSPA1A-GFP channel (cyan). **(B)** WGA-FA555 plasma membrane channel (magenta). **(C)** DAPI nuclear channel (blue). **(D)** Merged overlay of all three channels. Rows correspond to timepoints (37°C, 0h recovery, 8h recovery) and columns correspond to treatments (DMSO, Exo1, Exo2, BFA, Tun) as in Fig. 2A. Scale bar = 10 μm.

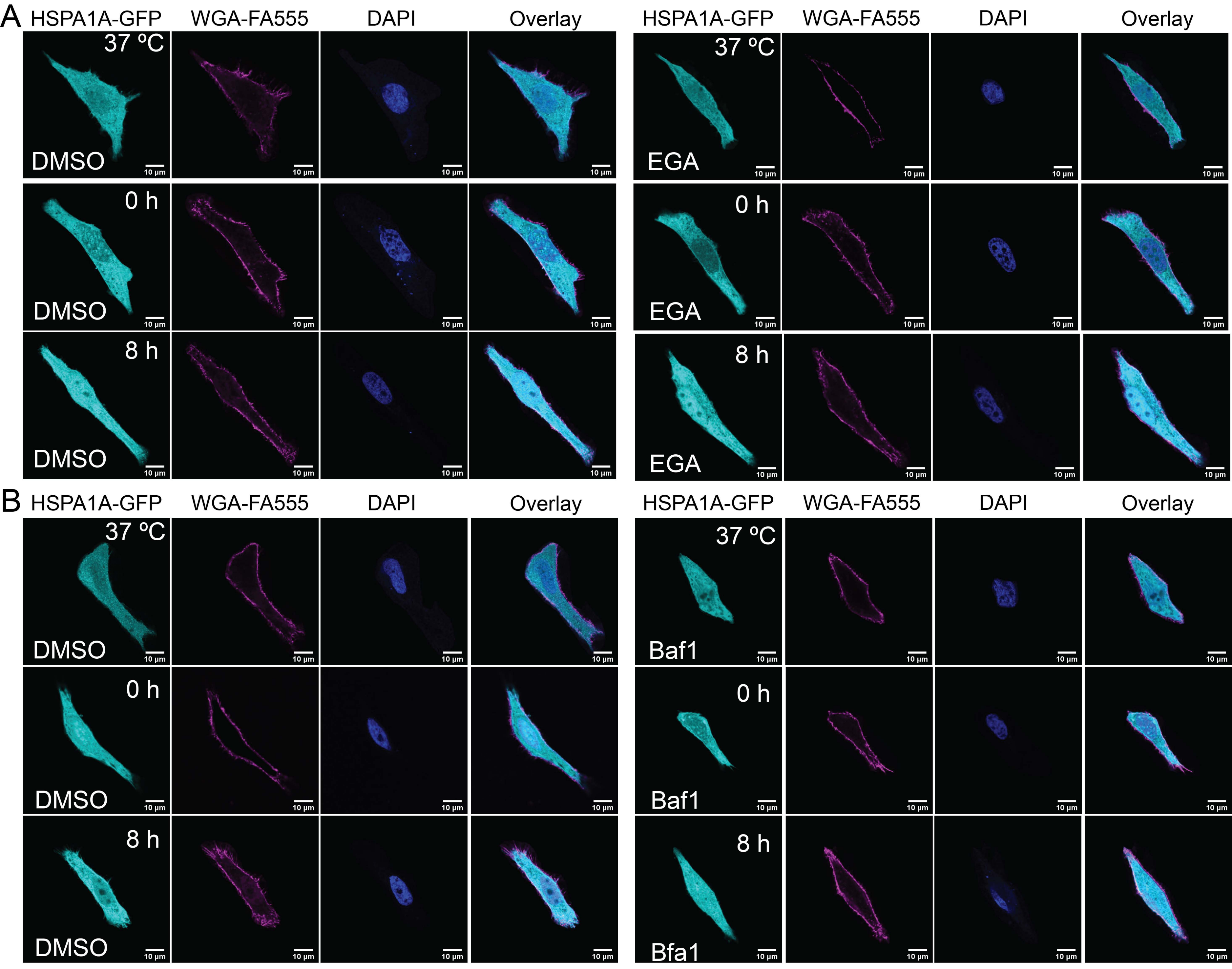
A
HSPA1A-GFP
WGA-FA555
DAPI
Overlay
37 ºC
DMSO
0 h
8 h
EGA
B
Baf1
Bfa1
10 µm

**Supplemental Fig. S3. Individual channel images corresponding to Figure 3A and 3C.** HeLa cells expressing HSPA1A-GFP were treated with DMSO, EGA, or Bafilomycin A1 (BafA1) and maintained at 37°C or subjected to heat shock (42°C, 1h) followed by recovery at 37°C for 0h or 8h. For each condition, individual channels and merged overlay are shown: HSPA1A-GFP (cyan), WGA-FA555 plasma membrane marker (magenta), and DAPI nuclear stain (blue). **(A)** DMSO and EGA conditions corresponding to Fig. 3A. **(B)** DMSO and BafA1 conditions corresponding to Fig. 3C. Rows correspond to timepoints (37°C, 0h recovery, 8h recovery). Scale bar = 10 μm.

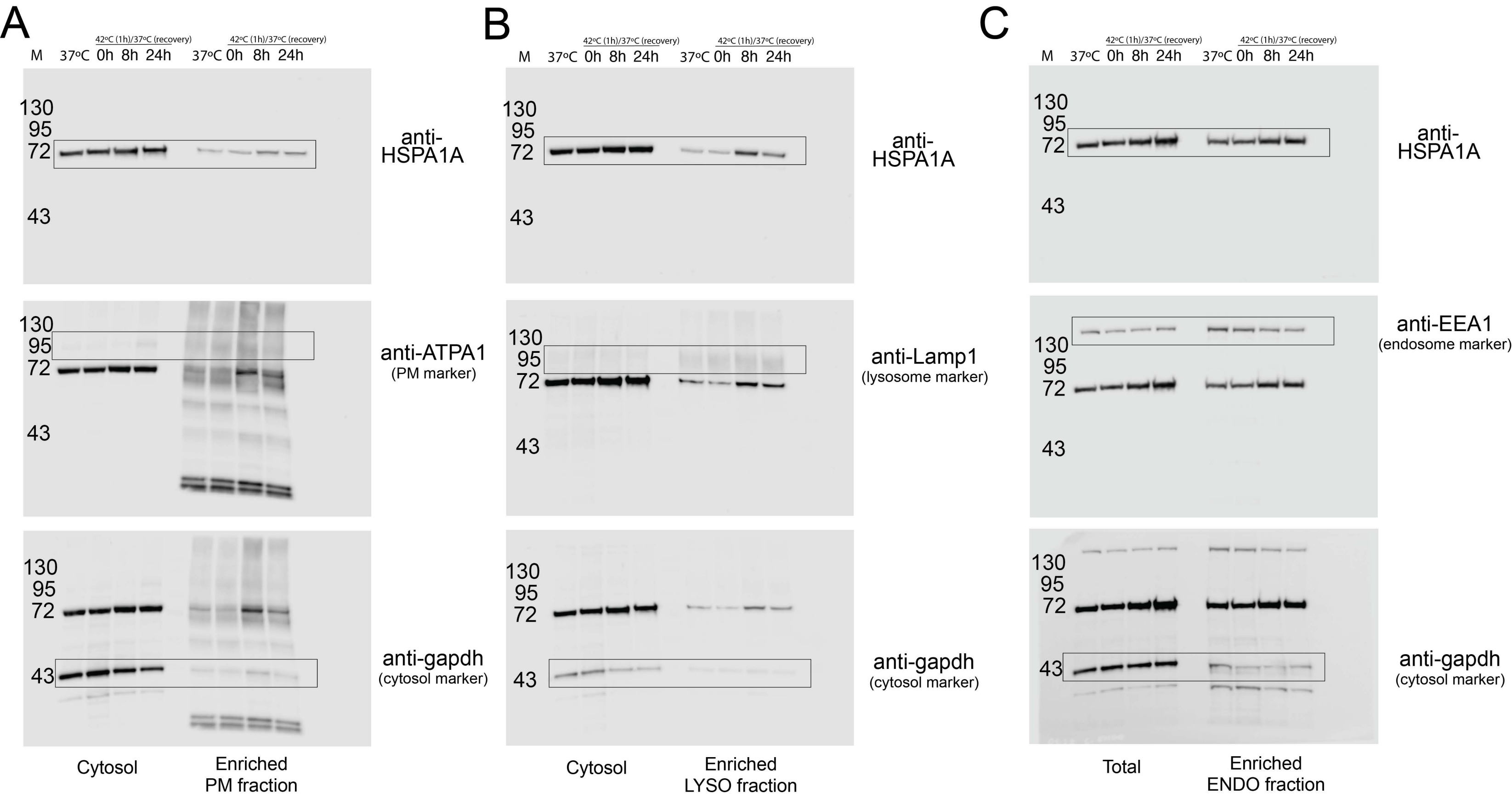
A
B
C
M
37°C
0h
8h
24h
42°C (1h)/37°C (recovery)
130
95
72
43
anti-HSPA1A
anti-ATPA1
(PM marker)
anti-gapdh
(cytosol marker)
anti-Lamp1
(lysosome marker)
anti-EEA1
(endosome marker)
Cytosol
Enriched PM fraction
Enriched LYSO fraction
Total
Enriched ENDO fraction

**Supplemental Figure 4. Subcellular fractionation western blot analysis of HSPA1A redistribution following heat shock.** HeLa cells were maintained at 37°C or subjected to heat shock (42°C, 1h) and collected at 37°C, 0h, 8h, and 24h recovery. Subcellular fractions were isolated and analyzed by western blotting. **(A)** Cytosolic and PM fractions probed with anti-HSPA1A, anti-ATP1A1 (PM marker), and anti-GAPDH (cytosolic marker). **(B)** Lysosomal fractions probed with anti-HSPA1A, anti-LAMP1 (lysosomal marker), and anti-GAPDH (cytosolic marker). **(C)** Early endosomal fractions probed with anti-HSPA1A, anti-EEA1 (endosomal marker), and anti-GAPDH (cytosolic marker). M, molecular weight marker; sizes indicated in kDa. Data represent a single biological experiment shown as qualitative biochemical reference consistent with the imaging-based redistribution patterns shown in Figure 1.

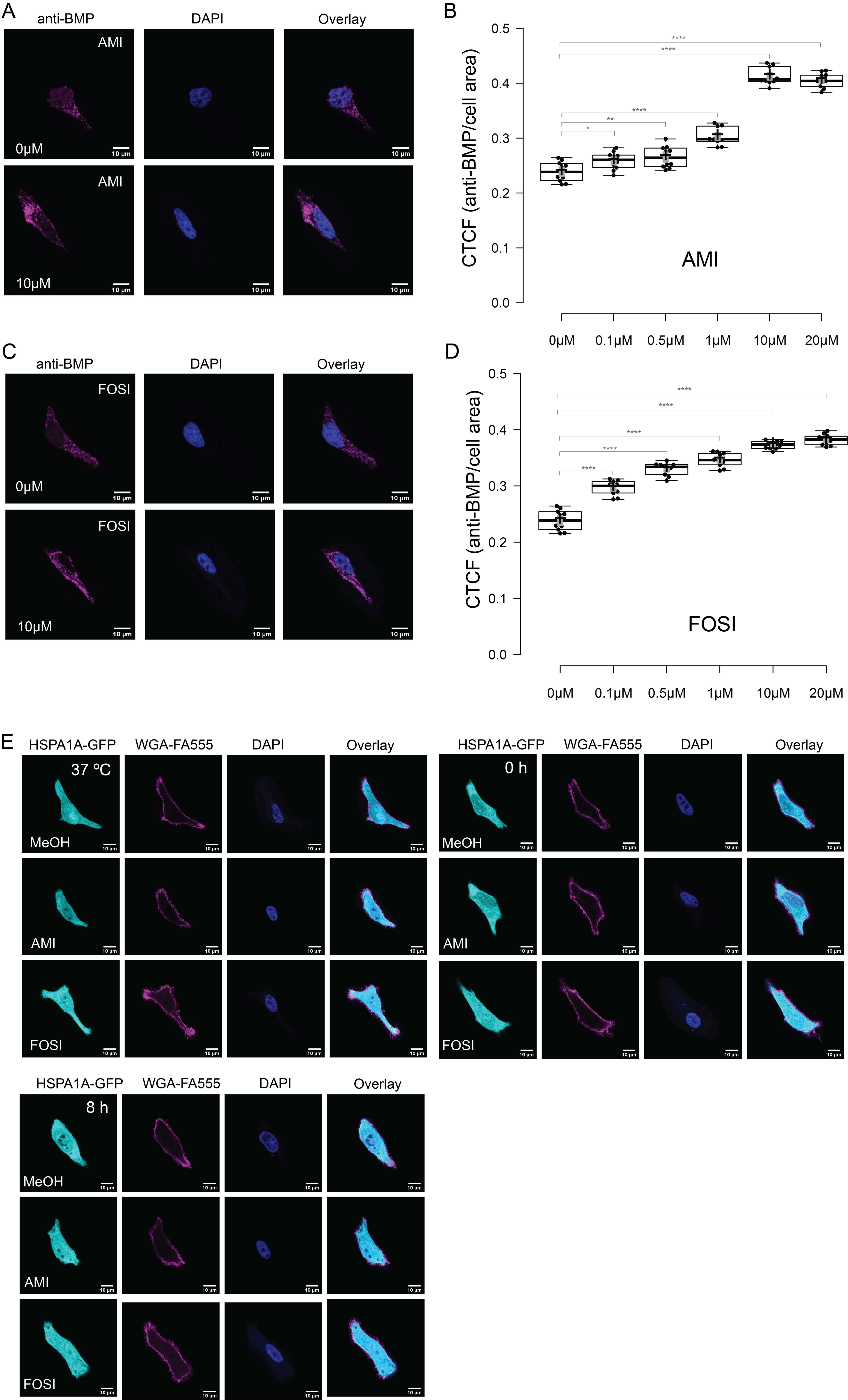

A
anti-BMP
DAPI
Overlay
AMI
0μM
10μM
10 μm
B
CTCF (anti-BMP/cell area)
0.0
0.1
0.2
0.3
0.4
0.5
AMI
0μM
0.1μM
0.5μM
1μM
10μM
20μM
C
anti-BMP
DAPI
Overlay
FOSI
0μM
10μM
10 μm
D
CTCF (anti-BMP/cell area)
0.0
0.1
0.2
0.3
0.4
0.5
FOSI
0μM
0.1μM
0.5μM
1μM
10μM
20μM
E
HSPA1A-GFP
WGA-FA555
DAPI
Overlay
37 ºC
0 h
8 h
MeOH
AMI
FOSI
10 μm

**Supplemental Fig. S5. Pharmacological validation of BMP modulation and representative images corresponding to Figure 4C and 4D.** HeLa cells were maintained at 37ºC and treated with increasing concentrations of Amiodarone (AMI) or Fosinopril (FOSI) for 24h. **(A** and **C)** Representative ICC images showing intracellular BMP immunoreactivity using anti-BMP antibody (clone 6C4; magenta) and DAPI (blue), with merged overlay only for the 0μM and 10μM conditions. Scale bar = 10 μm. **(B** and **D)** Dose-response quantification of intracellular BMP immunoreactivity by ICC following Amiodarone and Fosinopril treatment at increasing concentrations (0–20μM) for 24h at 37°C. Both compounds increased lysosomal BMP content in a dose-dependent manner. Amiodarone produced significant increases from 0.1μM ($p=0.020$) through 20μM ($p<0.0001$), with a plateau between 10 and 20μM. Fosinopril produced significant increases at all concentrations tested, from 0.1μM ($p<0.0001$) through 20μM ($p<0.0001$), without evidence of plateau. Statistical comparisons vs. 0μM baseline by unpaired t-test. * $p<0.05$, ** $p<0.01$, **** $p<0.0001$. n=20 cells per condition from three independent experiments. **(E)** Individual channel images for Amiodarone, Fosinopril, and MeOH vehicle control conditions corresponding to Fig. 4C and 4D, showing HSPA1A-GFP (cyan), WGA-FA555 plasma membrane marker (magenta), and DAPI nuclear stain (blue) at 37°C, 0h recovery, and 8h recovery. Scale bar = 10 μm.

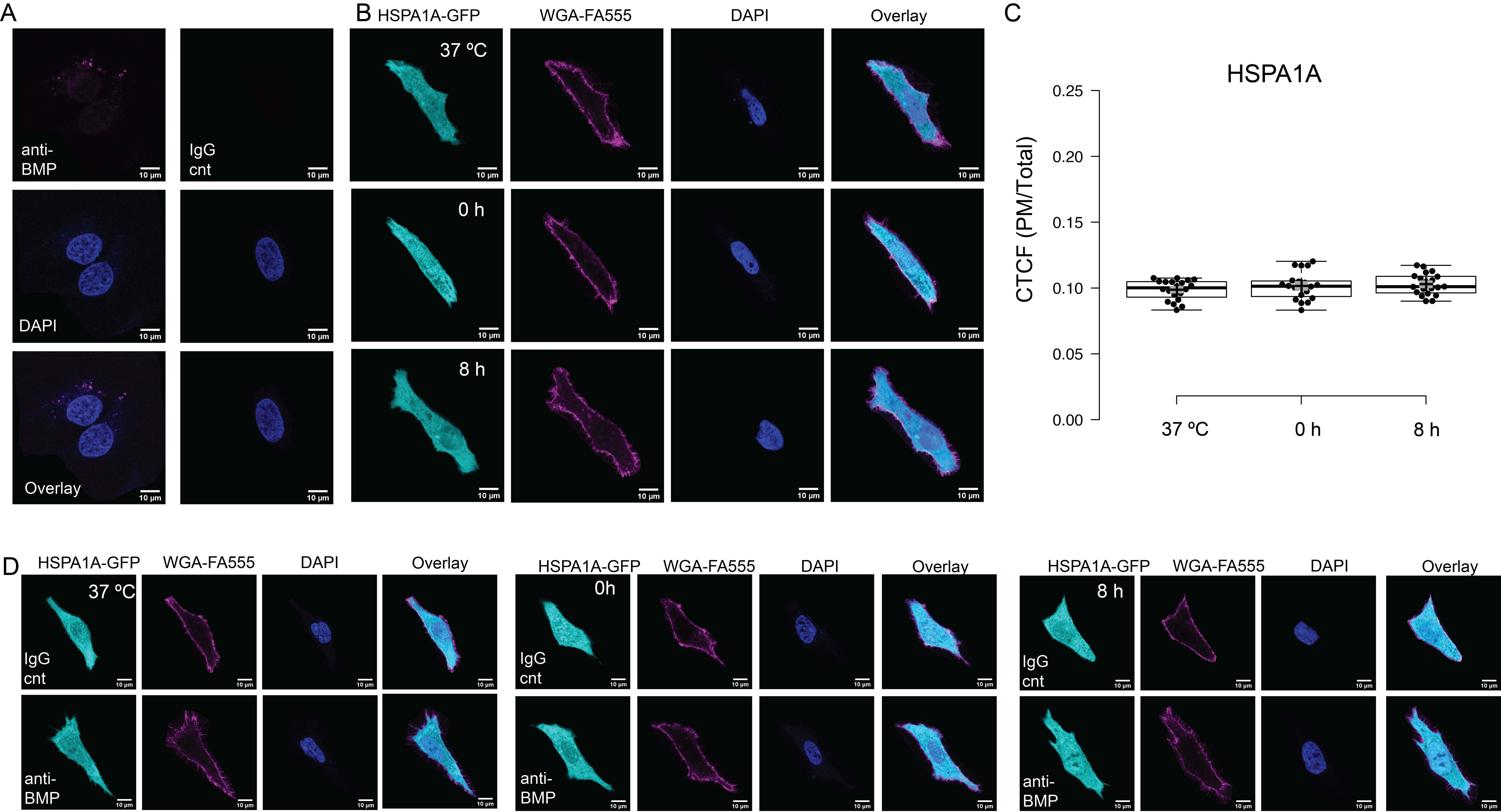
A
anti-
BMP
IgG
cnt
DAPI
Overlay
10 µm
B
HSPA1A-GFP
WGA-FA555
DAPI
Overlay
37 ºC
0 h
8 h
C
HSPA1A
CTCF (PM/Total)
0.25
0.20
0.15
0.10
0.05
0.00
37 ºC
0 h
8 h
D
HSPA1A-GFP
WGA-FA555
DAPI
Overlay
37 ºC
0h
8 h
IgG
cnt
anti-
BMP

**Supplemental Fig. S6. BioPORTER delivery validation and representative images corresponding to Figure 4E and 4F. (A)** Confirmation of intracellular antibody delivery by BioPORTER reagent, showing representative images of cells delivered anti-BMP antibody (clone 6C4; magenta) or mouse IgG isotype control (catalog #10400C; Invitrogen, Carlsbad, CA) at 37°C. **(B)** Representative confocal images of HeLa cells expressing HSPA1A-GFP delivered anti-PS antibody (anti-phosphatidylserine, clone 1H6; catalog #05-719; Sigma-Aldrich, St. Louis, MO) via BioPORTER and maintained at 37°C or subjected to heat shock (42°C, 1h) followed by recovery at 37°C for 0h or 8h. HSPA1A-GFP (cyan), WGA-FA555 (magenta), DAPI (blue). Scale bar = 10 μm. **(C)** Quantification of CTCF (PM/Total) for HSPA1A-GFP with anti-PS BioPORTER conditions shown in B. HSPA1A PM localization remained at baseline levels across all timepoints with no significant increase following heat shock (ANOVA $p=0.39$), confirming that BioPORTER delivers functional blocking antibodies capable of completely inhibiting lipid-dependent PM localization. Each data point represents one cell. Boxes show interquartile range with median; whiskers extend to 1.5× IQR. $n=20$ cells per condition from three independent experiments. **(D)** Individual channel images for anti-BMP and IgG isotype control conditions corresponding to Fig. 4E and 4F, showing HSPA1A-GFP (cyan), WGA-FA555 plasma membrane marker (magenta), and DAPI nuclear stain (blue) at 37°C, 0h recovery, and 8h recovery. Scale bar = 10 μm.

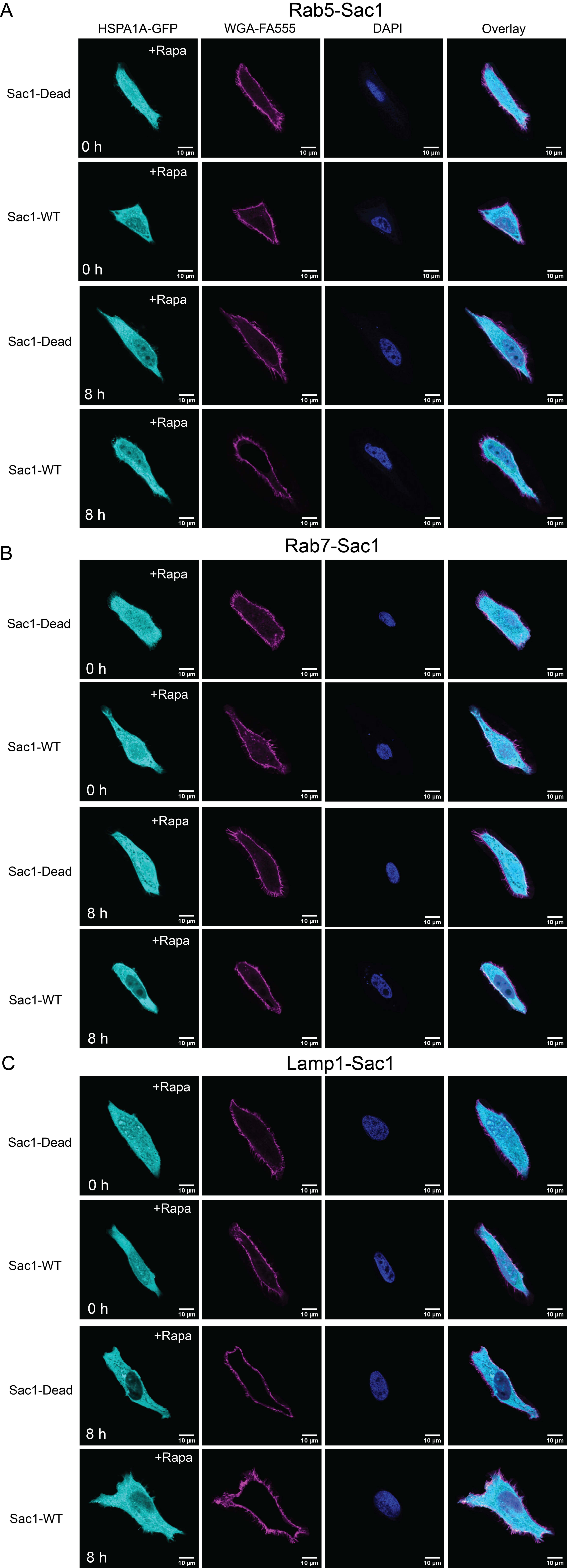
A
Rab5-Sac1
HSPA1A-GFP
WGA-FA555
DAPI
Overlay
Sac1-Dead
Sac1-WT
+Rapa
0 h
8 h
10 µm
B
Rab7-Sac1
C
Lamp1-Sac1

**Supplemental Fig. S7. Individual channel images corresponding to Figure 5.**
Representative confocal images showing individual channels for the key experimental conditions from Fig. 5. For each experiment, Sac1-Dead+Rapa and Sac1-WT+Rapa conditions are shown at 0h and 8h recovery following heat shock (42°C, 1h). Channels shown: HSPA1A-GFP (cyan), WGA-FA555 plasma membrane marker (magenta), and DAPI nuclear stain (blue), with merged overlay. **(A)** Rab5-targeted Sac1 recruitment to early endosomes. **(B)** Rab7-targeted Sac1 recruitment to late endosomes. **(C)** Lysosome-targeted Sac1. Scale bar = 10 μm.
